# Encoding of numerosity with robustness to object and scene identity in biologically inspired object recognition networks

**DOI:** 10.1101/2024.09.05.611433

**Authors:** Thomas Chapalain, Bertrand Thirion, Evelyn Eger

## Abstract

"Number sense", the ability to rapidly estimate object quantities in a visual scene without precise counting, is a crucial cognitive capacity found in humans and many other animals. Recent studies have identified artificial neurons tuned to numbers of items in biologically inspired vision models, even before training, and proposed these artificial neural networks as candidate models for the emergence of number sense in the brain. But real-world numerosity perception requires abstraction from the properties of individual objects and their contexts, unlike the simplified dot patterns used in previous studies. Using novel synthetically generated photorealistic stimuli, we show that deep convolutional neural networks optimized for object recognition encode information on approximate numerosity across diverse objects and scene types, which could be linearly read out from distributed activity patterns of later convolutional layers of different network architectures tested. In contrast, untrained networks with random weights failed to represent numerosity with abstractness to other visual properties, and instead captured mainly low-level visual features. Our findings emphasize the importance of using complex, naturalistic stimuli to investigate mechanisms of number sense in both biological and artificial systems, and suggest that the capacity of untrained networks to account for early life numerical abilities should be reassessed. They further point to a possible, so far underappreciated, contribution of the brain’s ventral visual pathway to representing numerosity with abstractness to other high-level visual properties.

## Introduction

Numerosity, or the number of objects in an environment, is an important parameter of variation in visual scenes that we can quickly apprehend in an approximate, ratio-dependent manner. In contrast to precise counting and the use of symbols based on language, this visual perceptual “number sense” is widespread across the animal kingdom (Lorenzi, Perrino, & Vallortigara, 2021; Nieder, 2021) and is present, at least in crude form, at very early stages of life (Izard, Sann, Spelke, & Streri, 2009; Xu & Spelke, 2000). Nevertheless, this basic nonverbal skill has been discussed as one possible foundational capacity enabling the development of more sophisticated numerical skills in humans (Halberda, Mazzocco, & Feigenson, 2008; Piazza et al., 2010; Schneider et al., 2017; Decarli et al., 2022), making its neural underpinnings a topic of wide interest.

Neuroscientific studies have provided evidence for a critical substrate underlying number sense predominantly in the dorsal visual pathway which leads up to the intraparietal cortex in the primate brain (Eger, 2016; Nieder & Dehaene, 2009). FMRI activity in these regions is sensitive to numerical changes in a ratio-dependent manner (Piazza, Izard, Pinel, Le Bihan, & Dehaene, 2004), and direct evoked activity patterns encode information about individual non-symbolic numerical quantities (Bulthé, De Smedt, & Op de Beeck, 2015; Castaldi, Piazza, Dehaene, Vignaud, & Eger, 2019; Cavdaroglu & Knops, 2018; Eger et al., 2009) with some independence from the sets’ non-numerical characteristics. Moreover, responses to specific numerosities are arranged in orderly topographic maps in the parietal cortex (Cai et al., 2021; Harvey, Klein, Petridou, & Dumoulin, 2013) with individual recording sites exhibiting Gaussian tuning curves to preferred numbers of items, similar to those observed in single neurons of macaque monkeys and other species (Ditz & Nieder, 2015; Kobylkov, Mayer, Zanon, & Vallortigara, 2022; Nieder & Miller, 2004; Viswanathan & Nieder, 2013).

Psychophysical work increasingly suggests that approximate numerosity is a perceptual property that is “directly sensed” within the visual system (Anobile, Cicchini, & Burr, 2016; Cicchini, Anobile, & Burr, 2016). Compatible with such a view, multiple studies have demonstrated that activity in early visual areas is already sensitive to the numerosity within visual displays (dot arrays) irrespective of a range of non-numerical properties such as the size and spacing of the items, e.g., (Park, DeWind, Woldorff, & Brannon, 2015; Fornaciai, Brannon, Woldorff, & Park, 2017; Fornaciai & Park, 2018; Castaldi et al., 2019; Paul, van Ackooij, Ten Cate, & Harvey, 2022). Nevertheless, a generally accepted computational account of how the brain extracts internal representations of numerosity from sensory responses to visual images is still lacking. While several computational studies within the field of numerical cognition have been dedicated to the problem of numerosity representation, some (Dehaene & Changeux, 1993; Hannagan, Nieder, Viswanathan, & Dehaene, 2017; Verguts & Fias, 2004) have not focused on explaining responses to visual images. Others have demonstrated that codes for numerical quantity can emerge from unsupervised learning in hierarchical generative networks (e.g., Stoianov & Zorzi, 2012; Testolin, Dolfi, Rochus, & Zorzi, 2020; Testolin, Zou, & McClelland, 2020). This work could explain aspects of behavioral and developmental findings related to numerosity processing, but remained limited to highly simplified stimuli and network architectures that bear little resemblance to the brain’s visual hierarchy.

In visual neuroscience more generally, hierarchical convolutional neural networks (HCNNs), a class of artificial neural networks used in computer vision that was initially inspired by aspects of visual neurophysiology, have emerged over the last decade as the mainstream models of high-level visual processing in the brain (Lindsay, 2020; Yamins & DiCarlo, 2016). These networks, which offer versatility of application to any type of visual image and allow for comparison of their internal representations with those found in the brain, have been particularly successful when applied to mechanisms of object recognition in the ventral visual stream (e.g., Yamins et al., 2014; Khaligh-Razavi & Kriegeskorte, 2014; Kubilius, Bracci, & Op de Beeck, 2016; Eickenberg, Gramfort, Varoquaux, & Thirion, 2017; Rajalingham et al., 2018).

In this context, an important recent finding was that individual artificial neurons in higher convolutional layers of such networks, after or even before training for object recognition, exhibited preferential responses to visual sets of variable numerosity very similar to those observed in biological neurons (Kim, Jang, Baek, Song, & Paik, 2021; Nasr, Viswanathan, & Nieder, 2019). Other work identified multi-scale spatial filters combined with normalization mechanisms as the critical ingredients for such numerosity information to arise (Park & Huber, 2022). This led to the suggestion that visual number sense could be an emergent property of suitable neural network architectures without dedicated neural mechanisms being optimized for numerosity identification per se. That number sense could emerge from the same neural machinery as the one that supports object recognition nevertheless seems counterintuitive: The notion of “four” applies in the same way to four cats as well as to four guitars, regardless of their precise visual properties and the context of their appearance. Thus, it requires generalization across the very same factors that visual systems need to be sensitive to in order to recognize objects. Given that previous studies investigating responses to numerosity in CNNs have only tested these networks with binary dot patterns (as is also typically done in cognitive neuroscience studies), it remains possible that the previous findings hold only for this specific case of simple contrast defined patterns. In such stimuli, it becomes difficult to dissociate responses to the discrete number of items per se from other factors that could be used as a proxy to numerosity in the networks. For example, for simple binary dot patterns with fixed luminance change between items and background, a larger number of items implies a difference in the spatial frequency distribution (Dakin, Tibber, Greenwood, Kingdom, & Morgan, 2011; Morgan, Raphael, Tibber, & Dakin, 2014). Moreover, recent work has shown that even across changes in non-numerical attributes such as the size and spacing of items, numerosity in dot patterns remains closely related to aspects of low-level contrast statistics expressed in terms of Fourier magnitude (Paul et al., 2022). This implies that different responses to variable numbers of items could arise from very simple sensory mechanisms that would unlikely be sufficient to extract numerosity in more general and naturalistic settings.

If artificial neural networks would fail to discriminate numerosity when they are tested with more complex and naturalistic stimuli that dissociate the number of discrete items from such low-level image statistics, then results such as those described might not reflect a sense of visual numerosity per se but simply responses to the correlated low-level statistics. If the sensory representations of networks optimized for object recognition were unable to support the simultaneous identification of objects and numerosity across changes in the objects’ characteristics, this could potentially explain why visual systems require separate resources for the two types of discrimination, and hence why numerosity processing has been found to be predominantly associated with the dorsal visual pathway in the primate brain. On the other hand, the finding that these two types of information can coexist in neural networks would provide hints towards a possible, and hitherto underappreciated, contribution of the ventral visual pathway to number sense.

Testing these possibilities requires stimuli that combine variability in high-level visual properties with experimental control. While recent work of others has shown that numerosity and non-numerical magnitudes can be extracted from large-scale datasets of natural images (Hou, Zorzi, & Testolin, 2024) opening the possibility for experiments using such stimuli, here we opted for a complementary approach involving original synthetic photorealistic stimuli that varied in object identity and scene context in addition to object number. This type of stimulus design allowed us to perform strong and well-controlled generalization tests that would not be easily feasible with fully natural real-world scenes for which some degree of association between object types, contexts and numbers are expected. It also helped us to dissociate the encoding of the discrete number of items per se from the encoding of several low-level visual summary statistics that we also quantified, which would be more correlated with numerosity in simpler stimuli or in paradigms that do not include similar generalization tests. Using this novel paradigm, we examined whether numerosity can be read out from the population activity of networks optimized for object recognition and their untrained counterparts, and tested for the degree to which such decoding can abstract from changes in object and background scene content. We also systematically investigated the extent to which numerosity decoding performance can be explained by several low-level summary statistics of the presented images. Finally, we examined numerosity-selective artificial neurons defined as in previous work for their ability to encode numerosity in our photorealistic stimuli, and compared results obtained with our photorealistic stimuli to those obtained with more simple binary patterns.

Our findings point to a population code for numbers of objects that supports some degree of independence from object and background changes, which is not carried by a few artificial neurons with the highest selectivity for binary dot patterns as those described in previous studies, and present only in object recognition trained (but not untrained) networks.

## Results

### Photorealistic stimuli decorrelate numerosity from low-level image statistics

We generated two stimulus datasets of synthetic photorealistic images comprising multiple objects and covering two different numerosity ranges. The “subitizing range” contained stimuli representing small numerosities (1, 2, 3 or 4 objects per image) and the “estimation range” contained larger numerosities (6, 10, 15 or 24 objects per image). For both datasets, each stimulus type is defined by the choice of an object, a background scene and a numerosity, and the chosen object is inserted into the background at randomly chosen positions and views. The choice to combine objects with unrelated background scenes is an adaptation of the strategy used with single objects in several influential studies investigating object representations in HCNNs and neural responses, e.g. (Yamins et al., 2014), aiming to obtain rich stimulus variability. In addition to numerosity, we varied the size and spacing of our objects within four levels each (Fig 1A). This design allowed us to test the extent to which CNN representations discriminate numerosity across changes in objects, backgrounds or both, allowing for generalization across superordinate categories and finer-grained individual variation.

**Fig 1.**
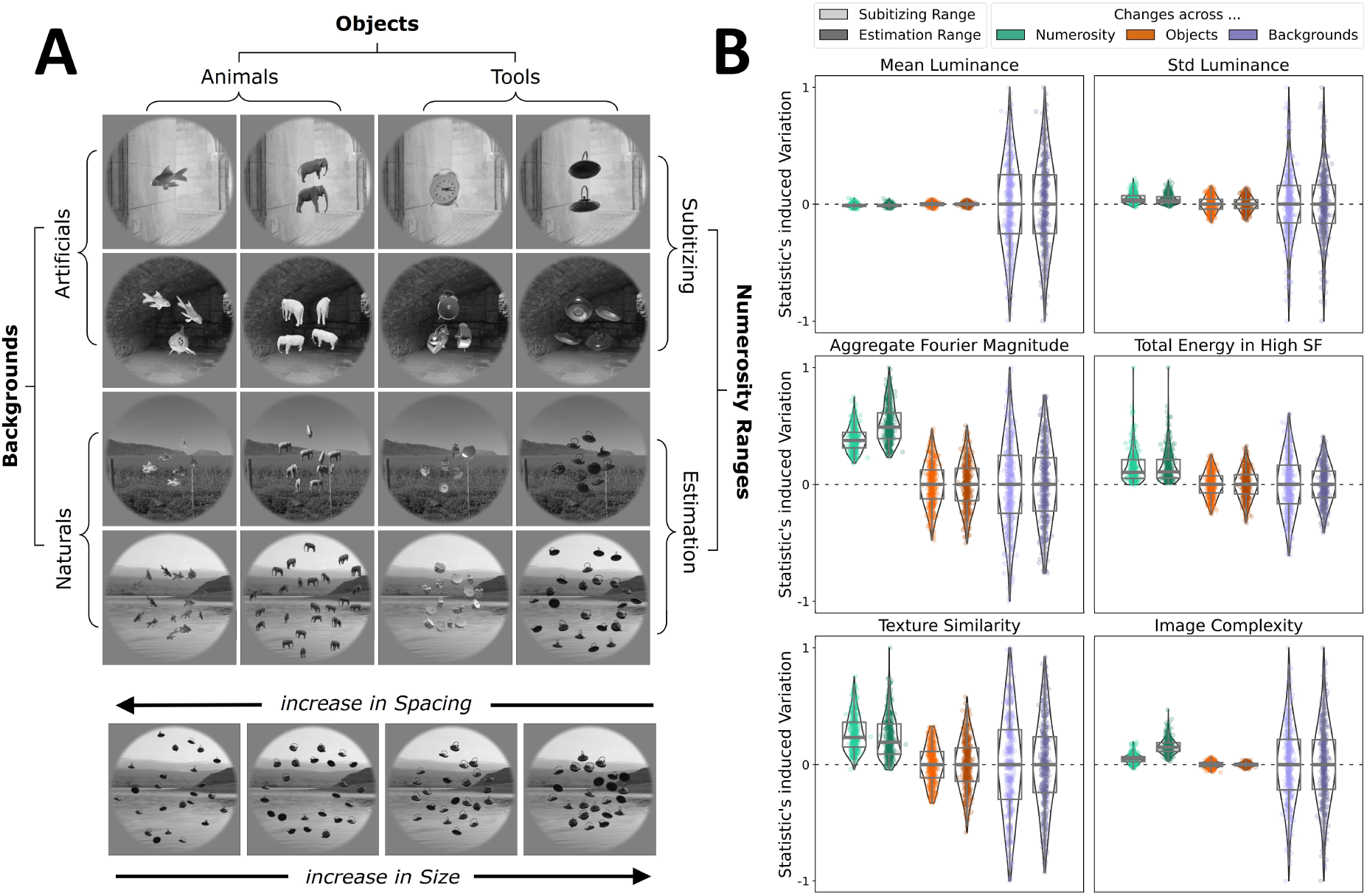
Stimulus design and quantification of image statistics. (**A**) Examples of synthetic photorealistic stimuli. The images vary in terms of object (two superordinate categories, animals or tools) and background scenes (two superordinate categories, natural or artificial). They contain different numbers of objects covering either small (subitizing: 1, 2, 3, 4) or large (estimation: 6, 10, 15, 24) numerical ranges. The objects are further varied in non-numerical quantities (4 levels of size and spacing) similar to previous work (DeWind et al., 2015). **(B)** Characterization of the stimulus dataset in terms of low-level summary statistics. Each panel shows how a specific low-level statistic varies when the stimulus numerosity (difference between largest and smallest), object or background (all pairwise differences) are changed. The variation is computed separately for each numerosity range (brighter or darker shade). For clarity, the plotted variation is normalized by its maximum absolute value. The whole distribution of each statistic’s variations is plotted as points, with the box indicating the median, first and third quartiles.

We characterized each stimulus image in terms of six different low-level statistics. The simplest ones correspond to luminance-related measures (mean and standard deviation of the image pixel values), which typically show some correlation with numerosity as long as there is also an increase in the total surface area of the objects, which is on average the case in our design. We also computed the energy at high spatial frequencies which has previously been proposed as a possible proxy for numerosity in dot sets (Dakin et al., 2011; Morgan et al., 2014) and could account for some psychophysical findings, as well as the aggregate Fourier magnitude as introduced by (Paul et al., 2022). Finally, we quantified two measures used in the field of natural scene representation (Scholte, Ghebreab, Waldorp, Smeulders, & Lamme, 2009) that describe the distribution of local contrast in natural images (referred to as “image complexity” and “texture similarity”, see Methods section for more details). In previous work these measures correlated strongly with activity in early visual cortex and we were interested in the extent to which they would provide a proxy for numerosity in photo-realistic images. Fig S1 and S2 provide an overview of our individual stimulus types in the space of these measures. The degree of variation in each measure when numerosity, object identity or background identity differed is summarized in Fig 1B. Although multiple statistics showed higher values on average for larger numerosities, changes in objects and backgrounds induced variations in the statistics that were in many cases as pronounced or even more pronounced (especially for background changes, see Table S1 for factors of increase in variation for objects and backgrounds with respect to numerosity). The Pearson correlation between numerosity and each of the 6 measures across all stimuli (computed separately for the subitizing / estimation range, respectively) was −0.02 / −0.01 for luminance mean, 0.08 / 0.07 for luminance standard deviation, 0.40 / 0.52 for aggregate Fourier magnitude, 0.23 / 0.28 for HSF energy, 0.08 / 0.23 for image complexity, and −0.26 / - 0.23 for texture similarity (for a more detailed overview of correlations split by either object or background, see Fig S3). These results suggest that by using analyses that leverage the high variability resulting from changes in objects and backgrounds on the one hand, and by comparing numerosity predictions from CNN representations with those based on the low-level statistics on the other hand, we can reveal whether there is a representation of numerosity information in CNNs that goes beyond these basic visual summary statistics.

### Numerosity information in CNNs generalizes across coarse object and scene categories

To determine the extent and nature of numerosity information in CNNs, we first investigated whether the networks’ feature representations allow for numerosity decoding with generalization across the broad semantic categories of the stimuli (animals vs tools, natural vs artificial backgrounds). Therefore, the internal representations of all stimuli were extracted from three common CNN architectures for object recognition, for five different convolutional layers of approximately equal relative depth across networks, here referred to as Conv1 to Conv5 (see Table S2 for details). This first step represents the encoding of our stimulus images (Fig 2A). We then performed a multivariate decoding analysis that splits the dataset into train/test sets separating the four superordinate categories (referred to as “coarse-grained generalization”, Fig 2B). This second step corresponds to the decoding of the logarithm of numerosity from the networks’ representations (Fig 2A). We found that for this type of generalization, only trained networks, but not their untrained counterparts, allowed for better than chance prediction of the number of objects in the image, with a clear gain in performance along the hierarchy of layers, from Conv1 to Conv5 (Fig 2C). The choice of using log(numerosity) rather than numerosity itself as target for decoding was made to account for the fact that numerosity discrimination is known to follow Weber’s law, but decoding with direct (linearly scaled) numerosity labels yielded qualitatively similar results (Fig S4).

**Fig 2.**
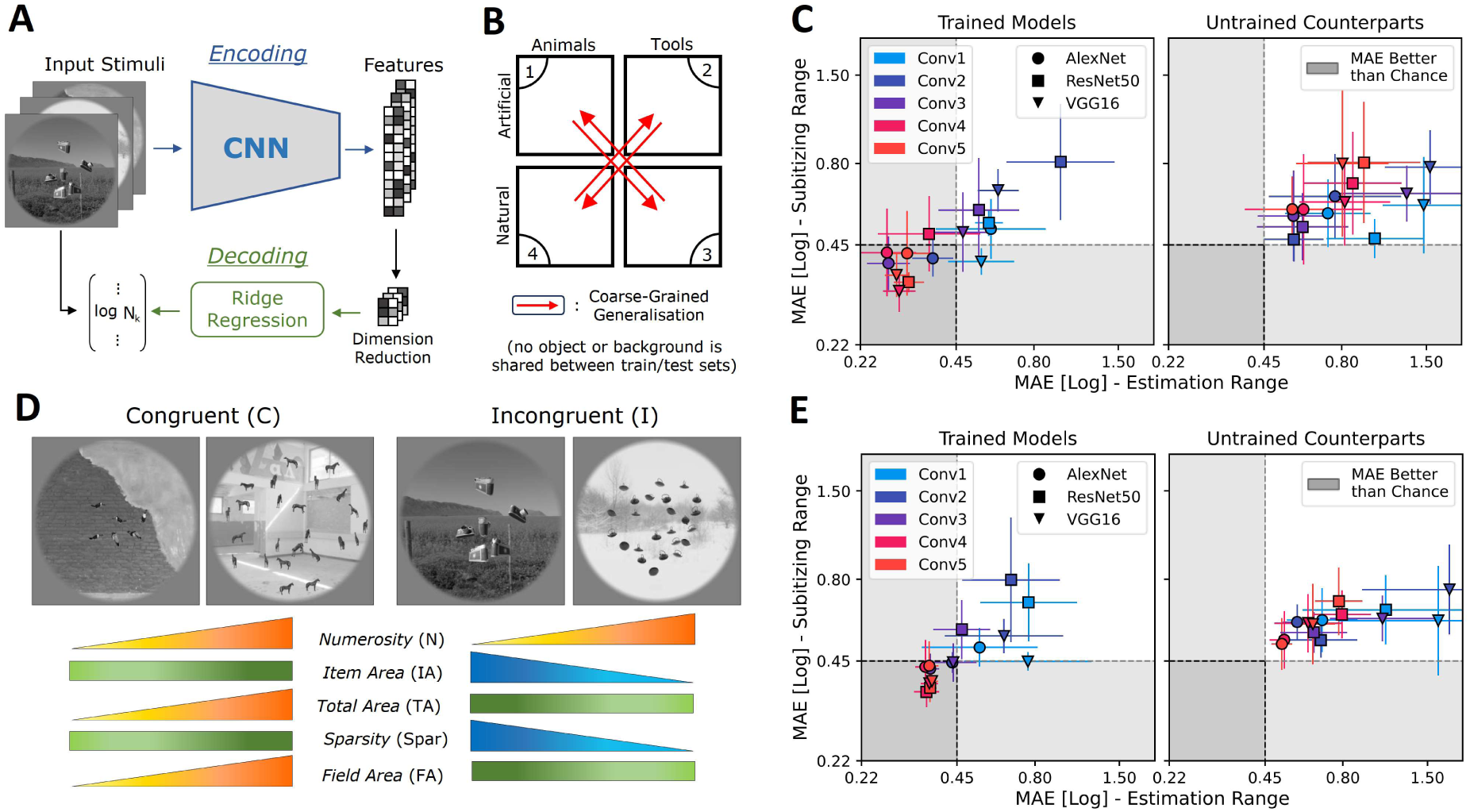
Numerosity information across coarse changes in visual properties. **(A**) Overview of analysis procedures. Encoding step: stimuli are fed to the CNN models. Their feature representations are extracted for several convolutional layers (Conv1 to Conv5) and undergo a feature selection. Decoding Step: Ridge regression is used to learn a linear mapping between the CNN representations of the stimuli and the logarithm of their numerosity. **(B)** Coarse-Grained generalization scheme: Decoders are trained and tested across different quadrants of the stimulus space corresponding to coarse semantic categories (red arrows). The performance is averaged for the predictive scores across all four train-test combinations. **(C)** Numerosity prediction performance (mean absolute error, MAE) in the coarse-grained generalization scheme for the different CNN architectures (markers) and layers (colors). Perfect performance is 0 and simulated chance performance is indicated by a dashed line. Error bars represent STD of the MAE over all the train-test combinations and cross-validation iterations. The subitizing range (y-axis) corresponds to 1 to 4 objects whereas the estimation range (x-axis) corresponds to 6 to 24 objects. **(D)** Stimulus subsets to control for effects of non-numerical quantities. Stimuli from each quadrant are divided into a congruent and an incongruent subset within which the non-numerical parameters (IA, TA, FA and Spar) are either increasing or decreasing with numerosity, respectively. Coarse-grained generalization is then tested between congruent and incongruent subsets. **(E)** numerosity decoding performance (MAE) for the different models on stimulus subsets from **(D)**, displayed as in **(C)**.

To check whether the lack of successful decoding from untrained networks could depend on the specific scheme used to randomly initialize the weights, we conducted additional analyses varying the hyperparameters of this initialization (see Methods for further details). We found that while these parameters had an influence on how much worse than chance the decoding from untrained networks performed, better than chance performance was not observed in any of the cases tested (Fig S5). These results point to the importance of network training for object recognition, as well as the hierarchical level of the layer representations, as two key factors facilitating the encoding of numerosity with some independence from high-level visual scene properties.

However, it is possible that the decoding of the number of objects in an image was driven by non-numerical quantities such as item or total surface area, sparsity or field area, which can never be decorrelated from numerosity all at the same time. In the way our stimulus space is designed, all four of these quantities had an equal degree of correlation with numerosity (Pearson correlation r = 0.35, see Fig S6). This suggests that the decoding analysis may have achieved better-than-chance predictive performance by using these quantities, rather than relying on a representation of numerosity per se.

To address this issue in a revised generalization scheme, in addition to the “coarse- grained generalization” split, we constrained the train and test stimuli so that their respective non-numerical features were made congruent (C) or incongruent (I) with numerosity (Fig 2D).

This meant that during decoder training, the item area (IA) and sparsity (Spar) of the stimuli remained constant, while their total area (TA) and field area (FA) increased with numerosity. In contrast, when the decoder was tested, the TA and FA of the stimuli were constant, while their IA and Spar decreased with increasing numerosity. With this additional constraint, networks trained on object recognition still allowed for decoding of numerosity across changes in high-level visual properties (Fig 2E), and as in the previous analysis, the predictive power for numerosity increased along the hierarchy of layers. Compared to the simple “coarse-grained generalization”, there was only a small decrease in decoding performance, with prediction errors increasing by on average 4% of the chance-level. These results show that the ability to read out numerosity from the internal activation patterns of CNNs cannot be mediated by the aforementioned non-numerical quantities. Taken together, these results suggest that when tested with our photorealistic stimuli, the approximate numerosity of objects is inherently encoded in object recognition trained networks, but not in their untrained counterparts, in a manner that is robust to coarse object and scene categories and classical non-numerical quantitative parameters.

### Generalization across exemplars and influence of low-level image statistics

To better understand across which image changes trained and untrained network representations can or cannot generalize, we investigated what was the degree of generalization of numerosity discrimination across individual objects and backgrounds, respectively. For each object (or background), we trained a decoder to estimate numerosity across all stimuli containing that specific object (or background). We then used the trained decoder to predict numerosity for each left-out object (or background), resulting in a pattern of pairwise generalization scores (Fig 3A).

When we applied this “fine-grained generalization” decoding analysis to the CNNs’ representations of the stimulus images, we observed a decrease in decoding performance for prediction of numerosity across changes in objects and backgrounds compared to testing on the same condition (Fig 3B). The cost in prediction performance with generalization was particularly pronounced for the untrained networks and in the case of background changes, which, according to our previous quantification of low-level image statistics, was the type of change that induced the most pronounced differences in these measures. The pattern of performance obtained for the untrained networks may therefore suggest that this decoding was driven by some combination of low-level image statistics rather than numerosity per se.

**Fig 3.**
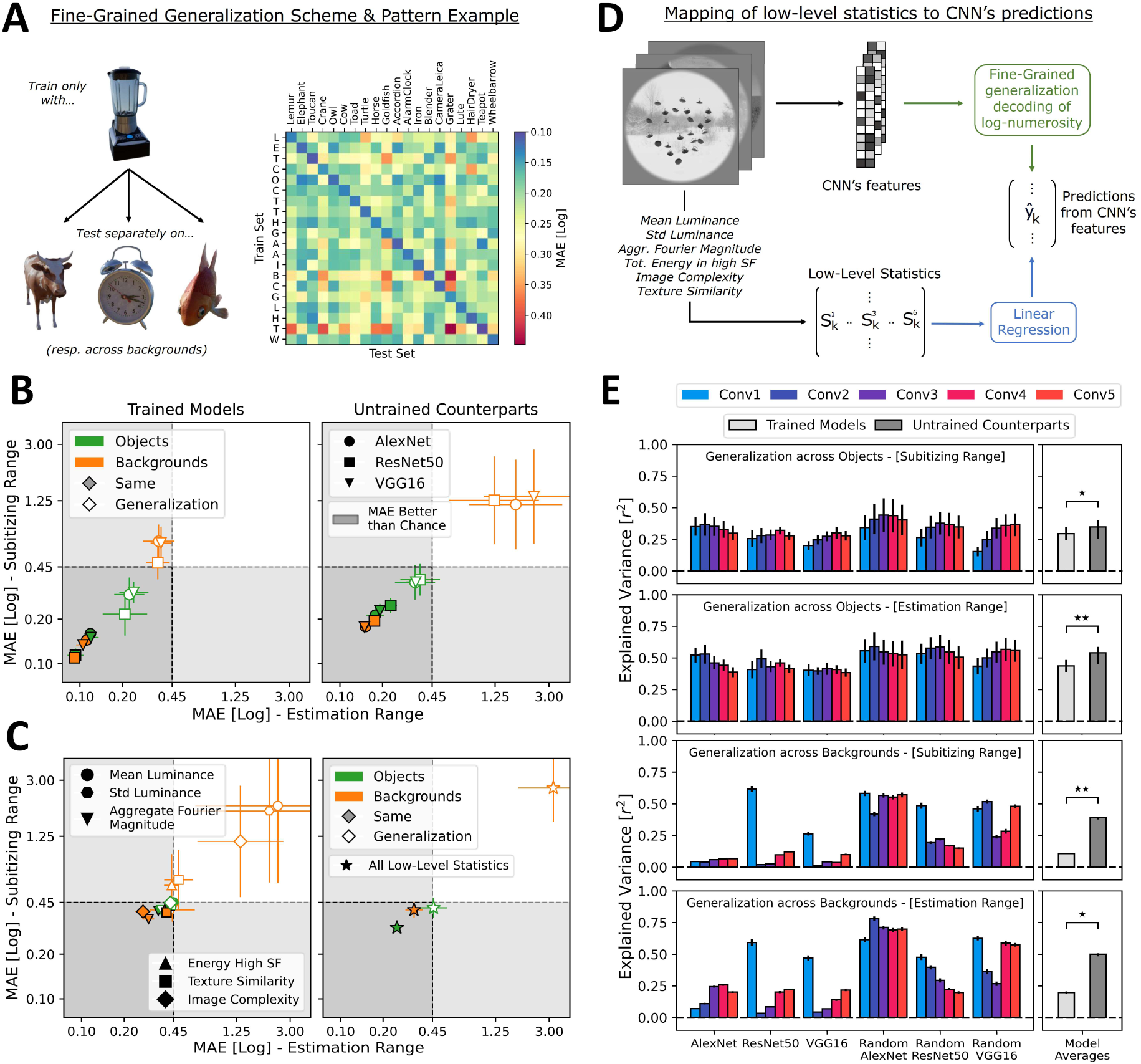
Numerosity decoding across fine changes in visual properties. **(A**) Fine-Grained generalization scheme: For each single object (or background), Ridge regression is trained to predict the number of objects across all stimuli containing this specific object (or background). The predictive performance is computed on each left-out object (or background) using MAE, leading to a pattern of generalization scores (an example of generalization across objects is given as a 20 × 20 matrix). **(B)** Numerosity prediction performance (MAE) for the different CNN architectures (marker shapes) for the fine-grained generalization across objects (green) or backgrounds (orange). Hollow markers represent performance across pairs of different objects (or backgrounds) and filled markers represent performance for test on the same condition (means and STD across all the train-test pairs). Performance is plotted for the most predictive layer of each model (see Table S3 for the corresponding layer labels). Perfect performance is 0 and simulated chance performance is indicated by a dashed line. The subitizing range (y-axis) corresponds to 1 to 4 objects whereas the estimation range (x-axis) corresponds to 6 to 24 objects. **(C)** Average numerosity decoding performance (MAE) of the Fine-Grained generalization for several simplistic models (marker shapes) using low-level image statistics as features for the Ridge regression. Decoding performance using all low-level statistics as a single concatenated input is shown in the right panel. **(D)** Analysis of effect of low-level statistics on predictions from CNNs: For every stimulus, six low-level statistics are computed and then mapped to numerosity predictions from the fine-grained generalization using multiple linear regression, yielding an explained variance score. **(E)** Contribution of the low-level statistics to numerosity predictions for the different CNN architectures and layers (colored bars), means and STD of the explained variance over all fine- grained generalization pairs. Explained variance averages across the trained (light gray) and untrained (dark gray) models are shown on the right. Significance levels for the difference correspond to Wilcoxon signed-rank tests: *p < 0.005, **p < 0.0005.

To further explore this idea of reliance on low-level image statistics, we performed two additional analyses. First, we tested what level of average performance could be achieved when trying to predict numerosity from the low-level statistics themselves. Using the six measures (luminance mean: L_m_, and standard deviation: L_sd_, high spatial frequency energy: E_high_, aggregate Fourier magnitude: M_agg_, image complexity: S_C_ and texture similarity: S_T_) as input features for multivariate decoding of numerosity, we found that they allowed for only marginally better than chance performance in a few cases where no generalization across objects and backgrounds was involved (Fig 3C). The numerosity predictions from the trained networks’ representations clearly and consistently outperformed the numerosity predictions from these low-level image statistics (both when each statistic was considered in isolation and when they were all used together). In contrast, the prediction performance from the untrained networks did not exceed that based on the low-level statistics by much. These results demonstrate that the generalization performance of trained networks cannot be explained by low-level image statistics alone, at least within the measures examined here. Conversely, the representations of untrained networks appear to function in a manner more similar to these simple low-level models.

Next, we reasoned that if the ability to decode numerosity from a network relied on some combination of the low-level image statistics as a shortcut, we would expect to see some relationship between the six image statistics and the numerosity predicted from the networks’ representations on a trial-by-trial basis. Therefore, we used multiple linear regression to test for an explicit linear mapping between low-level statistics and numerosity predictions on the test stimuli. This regression included the six image statistics together as predictors, and estimated their effect on the predictions obtained for generalization of numerosity decoding across individual objects or backgrounds. The explained variance computed from this analysis served as a metric to assess the extent to which the selected statistics could account for numerosity predictions across changes in stimuli (Fig 3D). We found that for both types of generalization, as well as for both numerosity ranges, the variance explained by low-level statistics differed significantly between trained and untrained networks, being higher for untrained networks (objects/subitizing: p = 1.16 × 10^−3^, objects/estimation: p = 6.10 × 10^−5^, backgrounds/subitizing: p = 3.05 × 10^−4^, backgrounds/estimation: p = 6.10 × 10^−4^, Wilcoxon signed-rank test). In particular, for generalization across backgrounds, the explained variance was nearly negligible for the trained networks, with the exception of Conv1 (Fig 3E). This layer has been documented in the literature to perform basic spatial filtering with weights similar to Gabor filters (Zeiler & Fergus, 2014), suggesting some degree of functional similarity to the hierarchical random filter banks in the untrained networks. These results further support the hypothesis that the number of discrete objects is encoded only in trained networks, whose numerosity prediction performance does not only clearly outperform the one of the low-level models, but also is less modulated by low-level statistics on a trial-by-trial basis. Numerosity decoding from the untrained networks instead substantially reflects basic visual summary statistics, which may account for the poor performance when generalizing across objects and backgrounds.

### The influence of non-numerical geometric quantities on numerosity predictions

While the previous analyses investigated the influence of several low-level image statistics on numerosity predictions, the degree to which these predictions could also be driven by some correlated non-numerical geometric quantities (such as total area, perimeter, density etc.) rather than numerosity per se is left open in these analyses. To address this issue, we took advantage of the structure of the parametric stimulus space which we had adapted from previously published studies (DeWind, Adams, Platt, & Brannon, 2015). Similar to this previous work, we estimated the influence of the three main orthogonal dimensions (Numerosity, Size in Area, Spacing) of the stimulus space on numerosity predictions by a multiple regression, to obtain insights into which of these and the further individual non-numerical quantitative dimensions the numerosity predictions were most aligned with. Results, summarized in Fig. 4 for all the networks tested, showed that while for trained networks predictions were roughly aligned with total perimeter when decoding from lower layers, they shifted to being closer to numerosity in higher layers. In untrained networks, on the other hand, predictions remained more aligned with total perimeter across all layers. An alternative visualization, showing the angle between the vector of estimated weights for Numerosity, Size in Area and Spacing and all the vectors representing the multiple individual axes of the stimulus space, is provided in Fig S7 for the example of network AlexNet.

**Fig 4.**
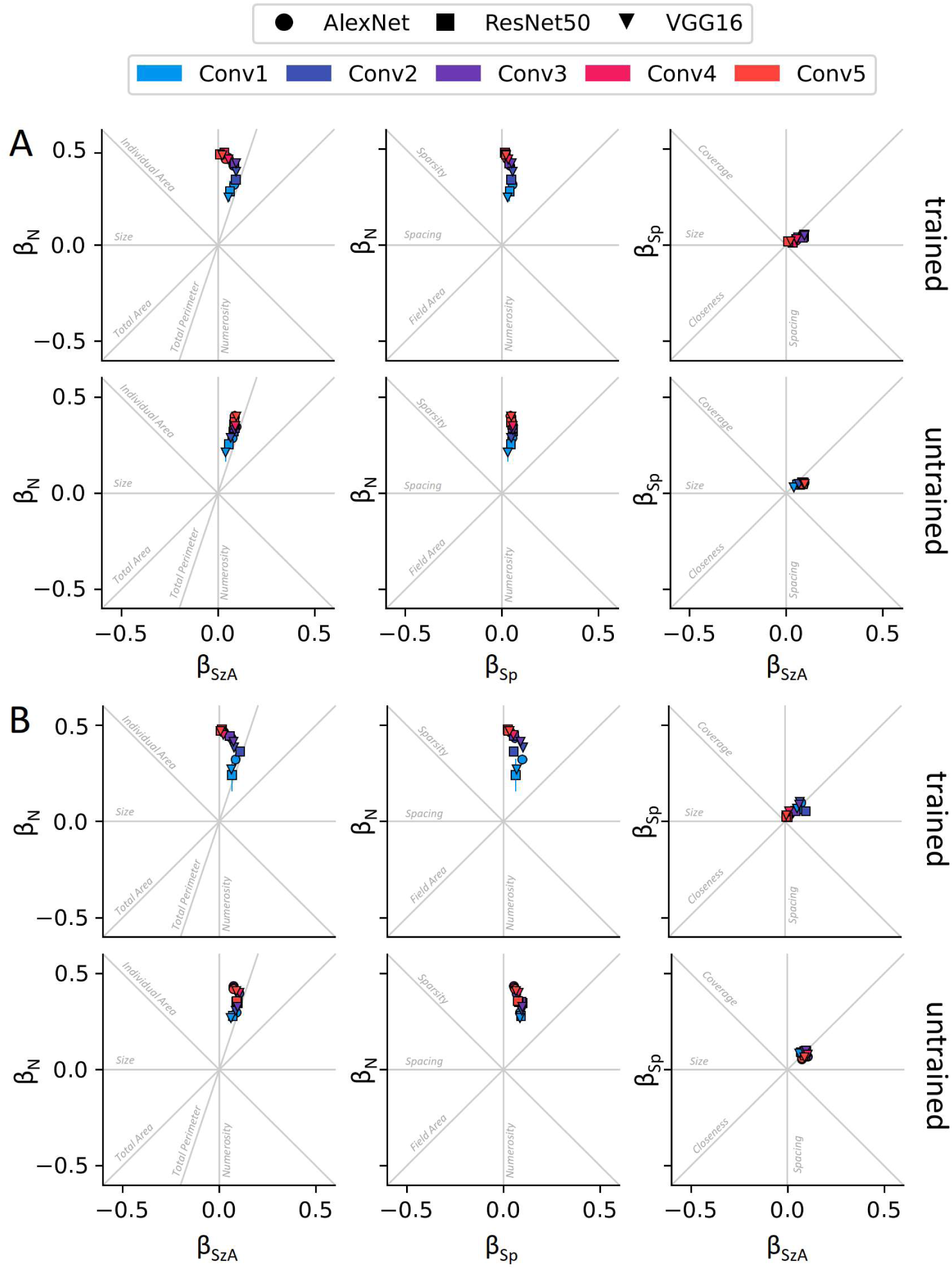
Numerosity prediction and effects of non-numerical quantity bias. Beta values (β_N_, β_SzA_, β_Sp_) resulting from a fit of numerosity predictions (for test on same object in the fine-grained generalization decoding analysis) by a multiple regression model including as predictors the three orthogonal axes of the parametric stimulus space (Numerosity, Size in Area, and Spacing), are plotted in the space of the quantitative dimensions, for all networks and layers. Panel A shows results for the subitizing range, and panel B for the estimation range. Markers represent means, and error bars STD of the estimated beta values across the 20 objects.

These results further reinforce our conclusion that it is the number of discrete items that is driving decoding performance from higher convolutional layers of object recognition trained CNNs. Non-numerical quantitative dimensions closely correlated with numerosity, such as total perimeter, can better account for decoding performance only in lower layers of trained networks, and in untrained networks overall.

### Simplified stimuli diminish differences between trained and untrained networks

Overall, in our investigation of the encoding of numerosity information in CNNs using synthetic photorealistic stimuli, we have observed important differences between object recognition trained and untrained networks that were not found in the previous literature. To confirm whether these differences were related to the use of more complex photorealistic stimuli, we next simplified our stimuli to determine the extent to which differences between the results of trained and untrained networks would vanish. To do this, we binarized each photorealistic stimulus image, setting the pixels enclosed by the objects to white, and background pixels to black. This resulted in a simplified stimulus set that more closely resembled the binary dot arrays used in previous studies. When the feature representations of the CNNs for these simplified stimuli were used as input for decoding of numerosity in the fine-grained generalization scheme across objects (Fig 5A), both trained and untrained networks showed better than chance decoding performance across all layers (Fig 5B). Testing for effects of model training (trained vs. untrained) and stimulus type (photorealistic vs. binary) in a 2-way repeated measures ANOVA revealed a significant interaction between training and stimulus type: (F(1, 29) = 368, p = 5.06 x 10^−18^), corresponding to a smaller difference between trained and untrained networks in the case of the simplified binary images compared to the photorealistic ones.

**Fig 5.**
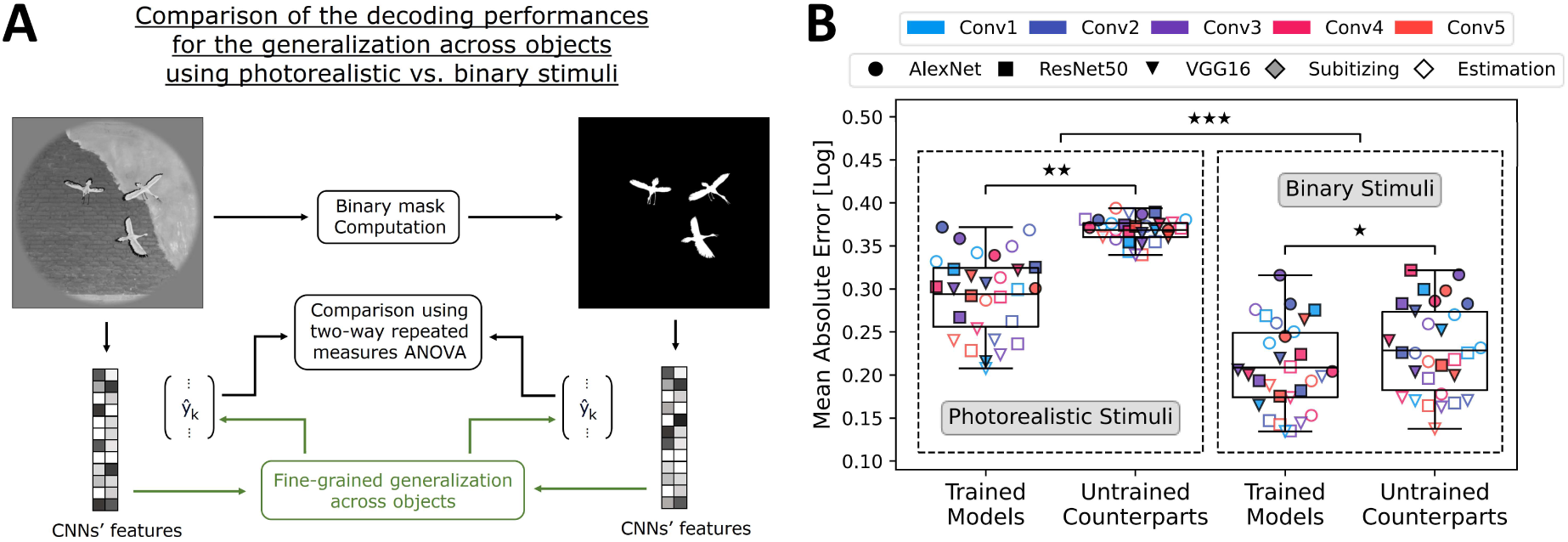
Impact of using photorealistic compared to simplified stimuli. **(A**) Overview of the analysis: A binarized version is computed from each original stimulus by setting pixels within objects to white and the background to black. Following this, the feature representation for this simplified stimulus dataset is extracted from all CNN models. **(B)** Comparison of the numerosity decoding performance (MAE) of the different CNN architectures (markers) and layers (colors) for the fine-grained generalization across objects, using the photorealistic or the simplified stimulus datasets. Filled and hollow markers represent decoding performance when using the stimuli from the subitizing and estimation range, respectively. The boxes extend from the first quartile to the third quartile of the data, with a line at the mean. Significance for the difference between decoding on photorealistic and simplified stimuli corresponds to the interaction in a two-way repeated measures ANOVA: ***p < 10^−15^. Significance for differences between trained and untrained networks correspond to Wilcoxon signed-rank tests: *p < 0.05, **p < 10^−5^.

While a discernible difference in performance between the trained networks and their untrained counterparts persisted in the case of the simplified stimuli (p = 0.016, Wilcoxon signed-rank test), the effect size was small: ΔMAE = 0.02, representing a relative difference of 9% from the average MAE across trained and untrained networks compared to 23% for the complex photorealistic stimuli (p = 1.73 x 10^−6^, Wilcoxon signed-rank test). These results show that the ability to extract numerosity from simplified stimuli such as binary sets of items, even with generalization across objects with pronounced shape changes, is not substantially different between untrained and trained object recognition networks, similar to the pattern of findings in the previous literature investigating dot-pattern tuned units. This further confirms that the crucial factor leading to differences between the results of object recognition trained and untrained networks here is the use of the photorealistic stimuli, rather than other ways in which our stimulus design or analysis differ from previous studies.

### Numerosity information in CNNs relies on a distributed population code

Given that previous studies (Kim et al., 2021; Nasr et al., 2019) reported artificial neurons discriminative of numerosity in AlexNet’s fifth convolutional layer (Conv5), with tuning characteristics that showed similarities to real neurons previously recorded in macaque monkeys, we wondered to what extent this type of neuron might underlie the encoding of numerosity in photorealistic stimuli as we found here. Therefore, we reproduced the analysis pipeline to identify these units. We generated the same types of dot arrays as used in the previous work, containing numerosities ranging from 1 to 30, in three stimulus sets which use different controls for non-numerical quantities such as dot size, area etc. We then tested the responses of AlexNet/Conv5 to these stimuli to find the set of artificial neurons that showed significant selectivity for stimuli with variable numbers of item (referred to as “number neurons” hereafter, see Methods for more details on their definition). We obtained N_s_=735 number neurons and N_ns_=997 non-selective neurons for AlexNet (1.70% and 2.3% of all units, respectively) and N_s_=233 and N_ns_=155 for random AlexNet (0.54% and 0.27% of all units, respectively) in the fifth convolutional layer (Fig 6A).

**Fig 6.**
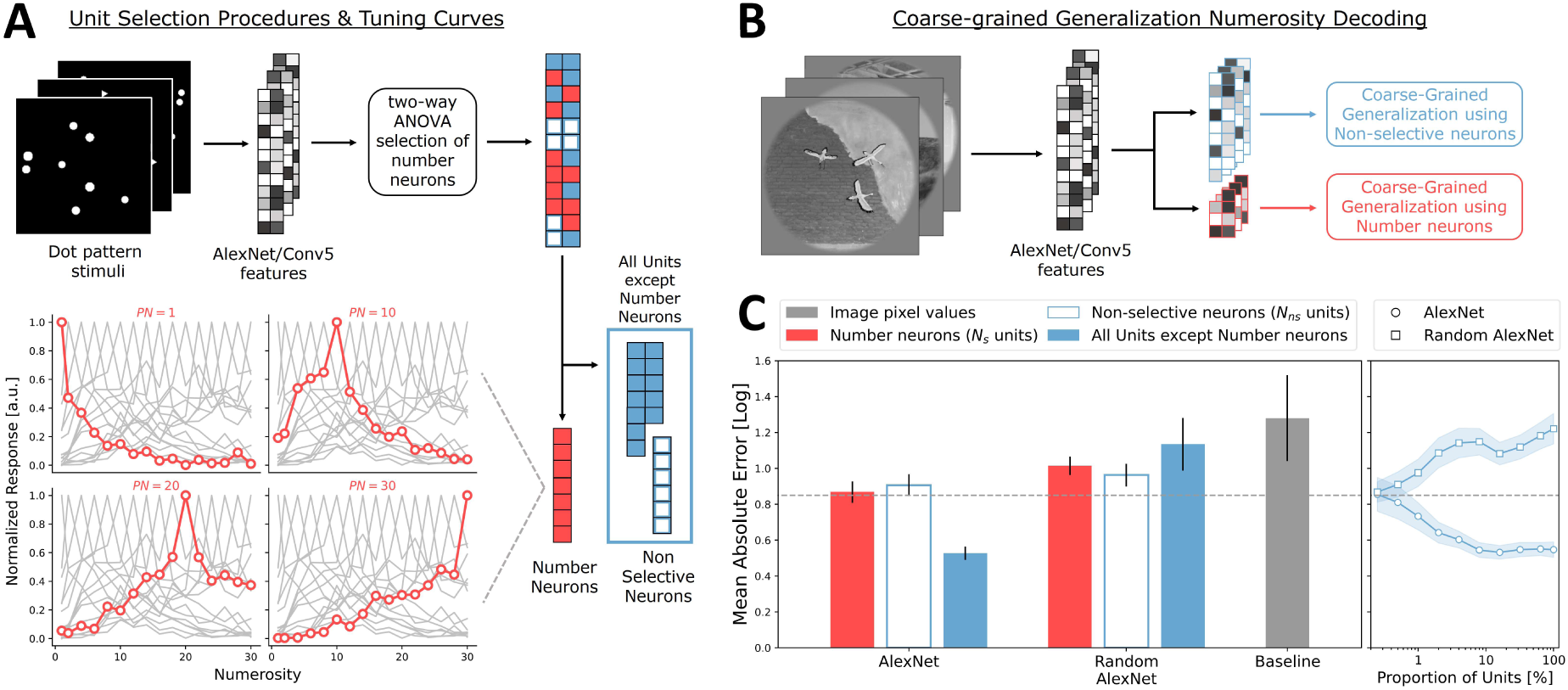
Analyses of numerosity selective units in dot patterns. **(A**) Analysis pipeline to identify number neurons: Using visual dot arrays similar to previous studies, we identified artificial neurons that exhibited significant selectivity to numerosity in AlexNet’s Conv5 (shown in red). Examples of the associated tuning curves for some preferred numerosities (PN) are also presented. Tuning curves are obtained by averaging the normalized activations of number neurons which share the same preferred numerosity. **(B)** Overview of the coarse-grained generalization using subsets of neurons: The set of AlexNet/Conv5 artificial neurons is split into the N_s_ number neurons (shown in red) and the N_ns_ non-selective neurons (shown in blue). Ridge regression decoders are trained and tested to predict the number of objects in photorealistic stimuli using each subset of neurons. **(C)** Numerosity prediction performance (MAE) for the fifth layer (Conv5) of AlexNet and its untrained counterpart (Random AlexNet) for the coarse-grained generalization (means and STD over all four train-test combinations). Perfect performance is 0 and chance performance is indicated by a dashed line. MAE scores for decoding based on the N_s_ number neurons and the N_ns_ non-selective neurons are shown in red and hollow blue, respectively. Decoding performance for the larger population excluding the N_s_ number neurons is shown in solid blue, and for the baseline of image pixel values in gray. MAE scores when randomly sampling different fractions of the entire artificial unit population are reported in the rightmost panel (means and std across all bootstrap iterations).

We then compared the ability of these number neurons to support decoding of numerosity with the one of non-selective neurons, for both AlexNet and its untrained counterpart, when using our previously described coarse-grained generalization approach on our photorealistic stimuli, this time on the union of the subitizing and the estimation range stimuli (Fig 6B). Using either the set of number neurons or the set of non-selective neurons, for both trained and untrained AlexNet, we observed a failure to generalize: decoding performance was either at chance or worse than chance, approaching the poor performance obtained when using image pixel values which served as a baseline (Fig 6C-left). However, we found that the larger population of artificial neurons, excluding the set of number neurons, allowed for successful decoding of numerosity for trained but not untrained AlexNet, replicating the results of the previous coarse-grained generalization analysis. This finding suggests that artificial neurons identified as being tuned to dot patterns are specific to such simplified stimuli and fail to consistently represent numerosity in the face of significant changes in high-level stimulus properties, such as changes in object and scene identity.

Finally, to further assess the influence of the number of artificial neurons used as input for decoding on the ability to read out numerosity, we repeated the previous type of decoding analysis while randomly sampling decreasing fractions of the total population of artificial neurons. We could distinguish two different regimes: First, when the fraction of units used in the analysis was reduced from the whole population to one tenth of it, the prediction performance remained constant, suggesting a high redundancy in the shared information across units. Then, when further reducing the number of units used to 108 (0.25% of the size of the full population), we observed a drop in decoding performance to chance-level, indicating that very few randomly selected units of the trained AlexNet could not encode numerosity in a way that generalized across object and background category changes (Fig 6C-right). Taken together, these results suggest that for the stimuli studied here, the representation of object numerosity with some tolerance to other high-level image changes is not carried by the artificial neurons showing selectivity to numerosity in dot pattern stimuli as those used in previous studies.

It might be argued that the previous analysis, replicating the methods for identification of numerosity selective artificial neurons from previous studies, did not identify the precise neurons most relevant for our type of stimuli. The stimuli used for unit selection previously differ from our new ones not only in being binary rather than photorealistic, but also, for example, in the way non-numerical quantities covary with numerosity. To assess numerosity selectivity of individual units under more comparable conditions, we made use of the binary masks created from our stimuli in addition to their photorealistic counterparts. When defining numerosity selective units in AlexNet’s conv5 layer based on either of these stimulus types (see Methods for details), we found that the overall number of units found to be selective was higher for the photorealistic than the binary stimuli, and that a tendency for more units preferring the largest or smallest numerical values was present in both cases (Fig. S8). A quantification of the overlap of units showed that the precise units which are found to be selective do depend to a large extent on whether the network is trained or not, on the stimulus type (binary vs photorealistic), and even to some extent on which of the two halves (“versions” only differing in the randomly chosen positions and view of the objects) of the dataset was used (Fig. S9). Results for numerosity decoding based on either numerosity-selective units or the larger population excluding the selective units for each stimulus type (Fig. S10) showed that on the stimulus type that they were selected on, the larger population excluding the selective units generally performed as good as or better than the selective units for discriminating numerosity with generalization across object and/or background changes.

These additional results further confirm the idea that the ability to encode numerosity with some robustness to other high-level visual properties as object or scene identity is a property of distributed population activity (of neurons which in isolation show little numerosity discrimination) rather than of a few single neurons which are maximally sensitive to numerosity and insensitive to size and spacing-related information.

## Discussion

Using original photorealistic stimuli, in this study we investigated the ability of deep convolutional neural networks, the current dominant models of high-level visual processing in the primate brain, to represent numerosity. Specifically, we tested the extent to which such networks can encode information about the approximate number of objects with invariance to changes in object and scene identity, for both small (1 to 4) and large (6 to 24) sets of objects. We found similar results across several common network architectures but a dissociation between the networks as a function of training: networks optimized for object recognition allowed for the number of objects in a visual scene to be read out from distributed patterns of activity in their higher layers, across the aforementioned changes in high-level visual properties, and this readout reflected numerosity rather than other closely correlated geometric quantities. Randomly initialized untrained networks, however, showed limited ability to represent numerosity information with robustness to other visual properties, especially when tested across changes in background which implied the most pronounced changes in low-level attributes in our stimuli. The failure of the untrained networks’ representations to support this type of discrimination appeared to be related to the fact that their predictions were to a greater extent driven by a combination of low-level visual summary statistics of the images, and more closely aligned with total perimeter than numerosity per se.

Our finding of a clear difference between the ability of object recognition trained and untrained networks to represent numerosity contrasts with the findings of previous studies investigating responses to numerosity in simple visual stimuli such as dot patterns, which suggested that networks such as those used here contain artificial neurons that respond preferentially to different numbers of items, both before and after object recognition training (Kim et al., 2021; Nasr et al., 2019). Our findings show that, contrary to previous suggestions (Kim et al., 2021), the mere projection through a hierarchy of convolution and pooling operations, without optimized feature weights, does not yield abstract numerosity information but at best something that can be a useful proxy to numerosity under very simplified stimulus conditions. In terms of the feature sets and computations required to extract numerosity, previous studies have found that in combination with divisive normalization, simple center surround receptive fields can be sufficient to obtain sensitivity to numerosity irrespective of size and spacing in the case of dot sets, but slightly more complex Gabor receptive fields are needed to extract numerosity and account for perceptual effects in stimuli comprising elements of different orientation (Park & Huber, 2022; Croteau, Fornaciai, Huber, & Park, 2024). An extrapolation of these ideas predicts that even more complex features would be needed for the more complex stimuli used here. Our findings show that higher layers of object recognition CNNs can support the generalizations that we tested, however, what is the minimally sufficient set of features and computations necessary for this remains to be determined.

Contrasting with the previous studies (Kim et al., 2021; Nasr et al., 2019) which focused on single unit selectivity as the putative code for numerosity in CNNs, we find that the information used to distinguish numerosity with some degree of robustness to object and scene context changes is not localized to few artificial neurons that are most selective to numerosity. The precise individual neurons showing numerosity selectivity did in fact depend to a fair degree on the stimuli used to select them (photorealistic or binary), but for each of these cases, the larger population of neurons excluding the selective ones allowed for at least as good or better numerosity decoding across object and/or scene changes. Thus, it appears that the information supporting this kind of invariance is widely distributed throughout the population of artificial neurons.

Our work compared numerosity information in CNNs for small and larger numbers of objects. In human perception, enumeration of small sets of items is typically fact and error-free, a phenomenon referred to as subitizing, but enumeration of larger numerosities under conditions when counting is prevented is approximate and follows a Weber’s law signature (Revkin, Piazza, Izard, Cohen, & Dehaene, 2008; Anobile et al., 2016) as long as the items are not too cluttered. We found no equivalent of the distinction between these two regimes in the networks studied here, where prediction of (the logarithm of) numerosity followed a similar approximate signature, and if anything, appeared slightly less accurate in the subitizing range. Differences in human performance in the subitizing and estimation range have often been attributed to the operation of two types of cognitive systems, one dedicated to object individuation/tracking and one dedicated to the representation of large approximate magnitudes (Feigenson, Dehaene, & Spelke, 2004; Piazza, Fumarola, Chinello, & Melcher, 2011). Interestingly, studies using dual tasks that occupy subjects’ attention have found that without the availability of spatial attentional resources, numerosity discrimination performance does indeed become approximate over the entire range (Vetter, Butterworth, & Bahrami, 2008; Burr, Turi, & Anobile, 2010). Given that spatial attention involves recurrent/feedback processing that is not present in the purely feed- forward network architectures studied here, the present finding of approximate performance across the whole numerosity range is not surprising and is indeed consistent with the human literature.

The artificial vision community has long recognized that the representations learned by CNNs trained on object recognition can contain some information useful for other visual tasks as well (Donahue et al., 2013; Razavian, Azizpour, Sullivan, & Carlsson, 2014). Still, it may be somewhat surprising that numerosity is one of them, given that identifying objects as opposed to numerosity across changes in objects or other high-level visual properties, intuitively seems to present quite orthogonal computational challenges. However, the results obtained here for numerosity nicely complement some findings in the previous literature on the representation of objects and other types of category-orthogonal visual information: For the encoding of the size and location of individual objects, it has been observed that, even though these properties are orthogonal to the task objective of an object recognition CNN, information about them is not lost but is actually enhanced or encoded more explicitly along the hierarchy of network layers, as it is also along the visual hierarchy of the brain (Hong, Yamins, Majaj, & DiCarlo, 2016). In addition, information about object pose and background appears to be encoded in distributed population activity in a way that is “factorized” and thus separated from object identity (Lindsey & Issa, 2024). In these studies, distributed population codes achieved the greatest separation between the different types of information at later visual stages. This parallels our finding for numerosity, where the ability to read out this property across changes in object and scene category was also highest in later convolutional layers. The case of numerosity is nevertheless somewhat special given that variability in this property is not a prominent feature of the ImageNet data set used to train our models, which for the majority of images only contains a single object (Lin et al., 2015). Thus, while on the one hand our results provide further evidence showing that the representational capabilities of object recognition CNNs can be far richer than the types of discriminations on which these networks are explicitly trained, even when it comes to properties of sets of multiple objects, they also suggest that feature sets useful to distinguish numerosity are not a trivial consequence of having encountered multiple objects during training.

What kind of representational primitives could underlie the ability of the networks (or visual systems in general) to encode information about both what an object is and about how many objects there are independently of their detailed properties, without important trade-offs or even incompatibilities? While our own results cannot provide a definitive answer to this question, recent studies suggested that feature selectivity not only within object recognition CNNs, but also in the brain’s ventral visual pathway, is not specific to whole objects (Baker, Lu, Erlikhman, & Kellman, 2018; Jagadeesh & Gardner, 2022). More specifically, high-level visual representations at these stages have been found to be “texture-like” in that they can actually be “blind” to certain image changes that alter the global object shape (Jagadeesh & Gardner, 2022), contrary to what is typically assumed based on our perceptual experience of objects as whole entities. Rather than being a limitation, the authors suggested that these findings reflect the use of a general, flexible basis set of visual features that can provide versatility for many different types of visual discrimination.

Our results have important implications for the study of visual number sense in the human brain and behavior. First, they suggest that cognitive neuroscience research should not overlook a possible contribution of the ventral “vision for recognition” pathway to this ability. Most research in the domain of numerical cognition has focused on dorsal (fronto-parietal) areas, although some early findings in non-human primates suggested that a smaller percentage of number neurons also exist in the inferior temporal cortex (Nieder & Miller, 2004). The responses of these neurons, however, appeared to be less independent of non-numerical properties than those in fronto-parietal regions. More recently, similar neurons have also been reported in humans in the parahippocampal and entorhinal cortices (Kutter, Bostroem, Elger, Mormann, & Nieder, 2018), which can be considered as the highest levels of the ventral visual pathway. Moreover, recent evidence suggests that distributed fMRI activity patterns of the ventral pathway can encode information about the numerosity of dot sets independently of non-numerical (size and spacing related) parameters (Karami, Castaldi, Eger, & Piazza, 2023), as in previous reports for the dorsal pathway (Castaldi et al., 2019). While the numerical cognition field has made extensive use of dot sets as stimuli, few recent studies have used more naturalistic stimuli (Hofstetter & Dumoulin, 2022; Odic & Oppenheimer, 2023). Future studies should examine population activity in both ventral and dorsal pathways with stimuli and generalization tests similar to those used here, to understand the extent to which representations in both streams are equivalent or emphasizing complementary types of information, and which stages are most important in determining behavioral numerical discrimination performance in humans.

Individuals with pronounced difficulties in acquiring mathematical skills (in the most extreme cases referred to as developmental dyscalculia) (Butterworth, Varma, & Laurillard, 2011) often also show impairments in basic non-verbal number sense beyond symbolic numerical processing (Piazza et al., 2010; Schneider et al., 2017). Interestingly, these impairments have in some cases been found to vary with the degree to which the stimulus conditions require the separation of discrete numerosity from other quantities (Bugden & Ansari, 2016; Castaldi, Mirassou, Dehaene, Piazza, & Eger, 2018). Dyscalculics may also exhibit more general visual impairments related to the segregation and detection of visual objects, manifested by increased visual crowding, and reduced 2D shape and 3D structure from motion sensitivity (Castaldi, Turi, Cicchini, Gassama, & Eger, 2022; Castaldi, Turi, Gassama, Piazza, & Eger, 2020). When investigating visual number sense using dot sets, as is commonly done, a range of strategies based on low-level summary statistics rather than representations of segregated objects would be usable to achieve some success on the task. Therefore, re-examining non-symbolic number sense in dyscalculia with more complex and variable stimuli may make potential difficulties more apparent if they are due to impaired visual mechanisms for extracting multiple objects.

The presence of information on numerosity of dot pattern stimuli in untrained neural networks (Kim et al., 2021; Testolin, Zou, et al., 2020) has previously been proposed to explain how humans or other animals can already have some sensitivity to numerosity information very early after birth, which is subsequently refined during development (Piazza et al., 2010). It is commonly assumed that infants already possess relatively abstract numerical abilities, based on the fact that their numerical discrimination generalizes across sensory modalities (Gennari, Dehaene, Valera, & Dehaene-Lambertz, 2023; Izard et al., 2009). However, given that some low-level statistics closely follow numerosity in spatial dot sets (Paul et al., 2022), a similar relationship may exist between the number of flashes and equivalent measures in the temporal domain, and the question is to what extent generalization might occur at this level. At least some existing evidence suggests that the processing of spatial and temporal frequencies shows some correspondence between the visual and auditory modalities, respectively (Guzman-Martinez, Ortega, Grabowecky, Mossbridge, & Suzuki, 2012). Therefore, cross-modal sensitivity to numerosity in previous studies need not automatically imply that young infants’ visual systems would support the same type of invariant representation within the visual modality as we have tested in this study. Although some infant studies have used numerosity stimuli consisting of different objects (Strauss & Curtis, 1981; Starkey, Spelke, & Gelman, 1990; Izard, Dehaene- Lambertz, & Dehaene, 2008; Brez, Colombo, & Cohen, 2012), to our knowledge none have tested for generalization across both object and scene identities with important differences in low-level summary statistics, as we did here. If untrained networks are a sufficient model of newborns’ visual abilities, then we would expect that the ability to generalize numerical discrimination across similar types of changes would not be present very early in life. It is known that although basic visual function including some degree of shape discrimination is present soon after birth, the ability to discriminate object categories continues to mature up into adolescence, see (Ayzenberg & Behrmann, 2024) for a review. The possibility that the representation of numerosity with abstractness to other high-level visual changes would also show a somewhat protracted maturation related to other high-level visual abilities still remains open.

It is of course possible that the untrained CNN with completely random weights is an oversimplified and incomplete model of the newborn visual system. Mammalian visual systems do already exhibit preliminary tuning to at least some visual features at the onset of visual experience, which appears to develop under the influence of spontaneous organized waves of activity that occur in the visual pathways before birth or eye opening (Huberman, Feller, & Chapman, 2008). In addition, the brain’s subcortical visual pathways, which are thought to be precociously functional (Atkinson, 1984; Bronson, 1974), have no equivalent in the current neural network models and may be responsible for visual abilities going beyond those of the untrained networks studied here. In this context, it is of interest that some previous modeling work, which used deep belief networks to account for the developmental refinement of the behavioral precision of numerosity discrimination in dot pattern stimuli, found that a numerical acuity similar to the one of human infants could be obtained by partially untrained networks endowed with basic visuo-spatial filtering capabilities which might come closer to the preliminary visual feature selectivity present at birth (Zorzi & Testolin, 2017). More generally, it seems worth considering that visual number sense need not necessarily rely on a single mechanism, but that different computations and brain substrates could contribute in different ways across development. The fact that the relative salience of changes in numerosity, as opposed to object identity, has been found to change during early years of life (Brez et al., 2012) could be consistent with the involvement of multiple visual mechanisms that are active at different timepoints.

The numerical information observed here in artificial neural networks appeared to be widely distributed across the population of artificial neurons, but could be read out by a linear decoder trained to discriminate numerosity in a supervised fashion. This raises the question of how such information could be exploited in biological visual systems without the availability of explicit numerical labels, i.e., before learning to count. The fact that we have used a supervised decoder here does not necessarily mean that training with explicit numerical labels is strictly necessary for higher areas to be able to extract this information. Indeed, some recent modeling work interestingly suggests that a “number-line-like” representation of numerical magnitude can emerge at higher levels of a neural network after observing the changes in visual images when items are added, removed or just changed in position (Kondapaneni & Perona, 2024), at least for simplified dot sets. This suggests that feedback from action systems during spontaneous manipulation of objects and observation of the resulting visual changes could play an important role in the development or refinement of number sense.

The networks in which we observed numerosity information across changes in object and scene identity here were optimized for object recognition, which is the task that has made such networks widely popular in visual neuroscience research, and also the type of CNN that has previously been investigated in the context of research on numerosity representation. This does not mean that the findings obtained are necessarily specific to object recognition training per se, but it may be that numerical information in deep convolutional architectures arises in the context of experience with natural images in general, and is only modulated in degree by the demands of the trained task. An important goal for the future will be to investigate how the representations in object recognition networks differ from those resulting from optimization for other visual task goals, some of which might be hypothesized to be more relevant to the dorsal visual pathway’s general task goal of object-directed action. Comparing the representations of networks that differ in task goal and architecture in terms of their ability to account for brain representations in the dorsal and ventral visual pathways could provide important insights into the constraints that shape the emergence of number sense in the brain.

## Materials and methods

### Experimental design

The generation of stimuli for this study employed a set of twenty 3D object models (half animals, half tools) obtained from https://www.turbosquid.com/. Using Blender, a free and open-source 3D computer graphics software (https://www.blender.org/) with its Cycles rendering engine, photorealistic renderings of each 3D object model were generated from 48 viewpoints. These viewpoints are described via the standard unitary spherical coordinates (r, θ, ϕ) [deg], where the 3D model corresponds to the center of the sphere, θ ɛ {-45, -15, 15, 45}, and ϕ ɛ {+/-15, +/-45, +/-75, +/-105, +/-135, +/-165}. The reference angle (θ, ϕ) = (0, 0) was determined individually for each object. Specifically, these reference angles were established in an as far as possible semantically meaningful way, to ensure a reasonable degree of consistency across all objects. For instance, animals were consistently oriented with their heads upright and facing forward, while tools were positioned in alignment with how they are typically encountered in everyday life, with their primary rotational symmetry oriented along the vertical axis, if they possessed one. High Dynamic Range Images (HDRI) obtained from www.polyhaven.com were used to create the environmental background scenes, and to create naturalistic lighting for the objects. Each 3D object model was rendered with the lighting corresponding to 20 different scenes (half natural, half artificial), in addition to rendering a view of each background scene separately. Randomly drawn views of each given object were then inserted into each of the background scene renderings (see Fig 1), resulting in a total of 400 object-background combinations. The stimuli contained 1,2,3 or 4 objects for the small (subitizing) numerical range and 6,10,15 or 24 objects for the large (estimation) numerical range. The stimuli also varied in non-numerical quantities (4 levels of size and spacing, see Fig 1 and Fig S6 for additional details) similar to the approach used in previous work (DeWind et al., 2015). By creating two instances (with randomly chosen stimulus positions and views) for each set of parameters (numerosities, sizes, spacings, and object-background combinations), our photo-realistic stimulus dataset comprised a grand total of 51 200 stimuli for each of the two numerosity ranges (or 12 800 per numerosity).

The experiments were performed on grey-scale versions of the images given our aim of quantifying and testing effects of low-level image statistics (see below) which would be considerably less straightforward with added color contrast.

### Characterization of low-level image statistics

For all stimuli we quantified several low-level image statistics related to luminance, spatial frequency or local contrast content: Luminance represents the grayscale value, with L = 0 corresponding to black and L = 255 to a white pixel. We computed the mean of the luminance L_m_ and its standard deviation L_sd_ which represents the RMS contrast of the stimulus. By denoting L_ij_ the luminance of the (i, j) th pixel of a stimulus of size H × W, L_m_ and L_sd_ correspond to:

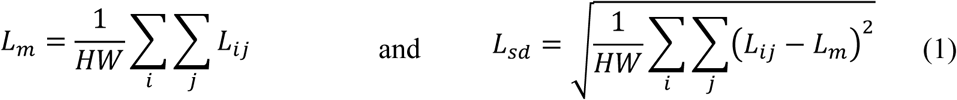

For the original 900-pixel images, the frequency range spans from ν_min_ = 1 cycle/image to ν_max_= 450 cycles/image. We computed E_high_, the total rotational average of the energy in the high spatial frequency range, by using a cut-off frequency (ν_cut_) between the low and high spatial frequencies. This cut-off frequency corresponded to a half-cycle matching the average object size: 184 pixels for the subitizing range and 75 pixels for the estimation range. By denoting F(ν_x_, ν_y_) the Fast Fourier Transform (FFT) of the stimulus at the spatial frequency (ν_x_, ν_y_) and E_r_(ν) its rotational average energy density spectrum at frequency ν, E_high_ can be expressed as:

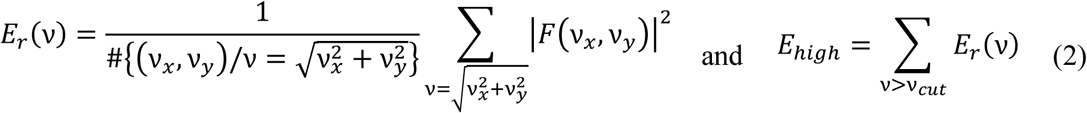

We also computed the aggregate rotational Fourier magnitude M_agg_, generalized from its definition in (Paul et al., 2022). M_agg_ corresponds to the sum of the Fourier magnitude at all orientations and spatial frequencies up to the first harmonic frequency ν_1_, which is a prominent characteristic of the FFT of a dot set. However, in the case of photo-realistic stimuli which use real views of 3D objects instead of dots, the boundaries of the harmonics do not manifest as clear throughs in the Fourier magnitude spectrum. In order to obtain a measure coming as close as possible to that used in the aforementioned previous work, we took advantage of the fact that ν_1_ depends on the size of the objects, as shown in (Paul et al., 2022), and we defined an equivalent cut-off frequency ṽ_1_that would correspond to the first harmonic frequency of a circle whose area is equal to the average area of the objects inserted in the photorealistic stimulus: The FFT of a circle of radius R is given by J_1_(νR)/ν where J_1_(.) is the first order Bessel function. If we denote IA_m_ the average area of the objects and R_e_ the equivalent radius of a circle covering the same area, a surrogate for this first harmonic frequency ṽ_1_ is given by:

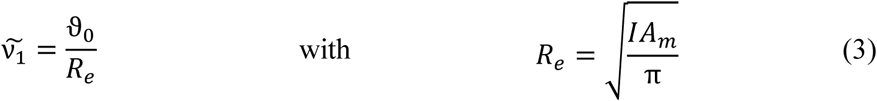

where ϑ_0_ corresponds to the first root of J_1_. A valid approximation of ϑ_0_ ≈ 3.83 can be found in numerical tables. Subsequently, the aggregate rotational Fourier magnitude M_agg_ can be derived as:

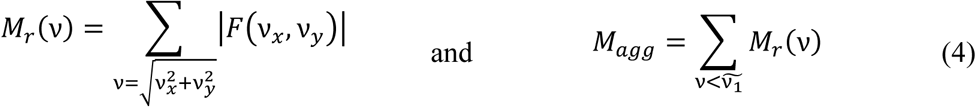

Measures of the “image complexity” S_C_ and the ”texture similarity” S_T_ were obtained from a Weibull fit of the local contrast distribution of our stimuli S similar to previous work (Scholte et al., 2009). We first obtained the local contrast C(i, j) at pixel (i, j) of the stimulus by filtering it with a derivative of a Gaussian kernel G_σ_, using a standard deviation of the Gaussian kernel σ = 12.

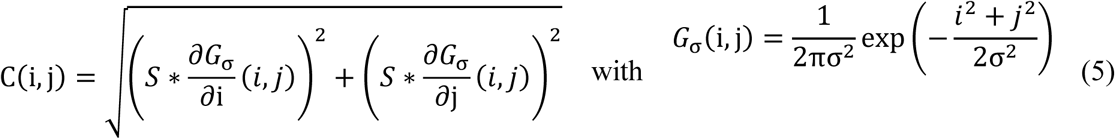

Then, we fitted the local contrast distribution C(c) with a Weibull distribution W of parameters (β, γ), using maximum likelihood estimation:

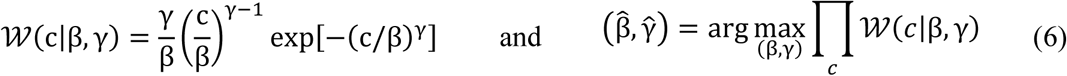

The stimulus representation of natural images in the space of the Weibull parameters (β,γ) is characterized by two main directions of variation, see also (Scholte et al., 2009). The first eigenvalue axis, corresponding to an increase in both β and γ, represents an increase in image complexity. The second eigenvalue axis shows a transition between images that have more or less homogeneous texture over the entire image. Accordingly, we obtain the “image complexity” and “texture similarity” measures of each stimulus by projecting its estimated 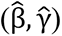 values onto the first and second eigenvalue axes of the Weibull parameter space of our entire stimulus set, respectively.

### Artificial neural networks and feature extraction

Three convolutional neural network (CNN) architectures commonly used for object classification were examined: AlexNet, ResNet50, and VGG16, all as implemented in PyTorch 2.0. (https://pytorch.org/). We used each of these CNNs in two instances: one pre-trained on the ImageNet dataset (Deng et al., 2009) and another untrained. The weights for the untrained networks were assigned using a LeCun initialization (LeCun, Bottou, Orr, & Müller, 2012), which corresponds to randomly sampling weights from a Gaussian distribution with mean 0 and standard deviation of 1/n_in (where n_in is the input size of the layer). To explore the influence of the set of hyperparameters of the random initialization on decoding performance, we also performed additional analyses that varied the standard deviation of the distribution by factors of of 1/500, 1/10, 2 and 10 with respect to the original LeCun initialization, and using either Gaussian or uniform distributions (Fig. S5). For our analysis, we focused on the post-ReLU activations of five convolutional layers located at comparable depth relative to the network’s size. A detailed overview of these layers is provided in Table S1. Stimulus images were down- sampled to the input resolution of the networks (224 x 224 pixels) and then fed to each of the selected networks to obtain their successive feature representations along the hierarchy of convolutional layers.

### Numerosity decoding analysis

We used linear Ridge regression as implemented in scikit-learn (https://scikit-learn.org/stable/) to predict the logarithm of numerosity from the CNN feature representations of the stimuli, and evaluated the decoding performance of each network on left-out stimuli using the mean absolute error (MAE) metric. Perfect decoding performance would result in a MAE score of 0. The optimal ridge penalization hyperparameter λ was selected by a nested leave-one-sample-out cross-validation loop during the training phase. To determine the chance level performance to be used as a baseline, the same Ridge regression was fit on a random permutation of the feature representations of the stimuli and their associated log-numerosity targets. In addition, to mitigate potential differences in statistical power arising from unequal dimensionality of the feature representations across networks and layers, we equated the dimension of each network’s feature space through univariate feature selection. Specifically, one-way ANOVA feature selection for any difference between numerosities was performed during the training phase, to reduce the dimension of each representation down to the smallest feature space size (i.e., 43 264 dimensions, AlexNet/Conv5) and subsequently applied during the testing phase.

In the coarse-grained generalization scheme, the stimulus dataset was split into train/test sets separating the four superordinate categories resulting in four subspaces, each consisting of 10 objects paired with 10 backgrounds (e.g., animals on natural backgrounds as training set and tools on artificial backgrounds as test set). For each of the 4 possible train-test set combinations, we performed a 6-fold cross-validation where in each cycle, we randomly selected 20 object/background conditions from the training set, resulting in a total of 1280 samples to learn the Ridge regression parameters. Within each fold, the MAE scores for this trained Ridge regression were then calculated using the entire test set. Finally, all the MAE scores were averaged across folds and the 4 train-test set combinations to obtain the cross-validated decoding performance score for a given network layer.

In the fine-grained generalization scheme, for each single object or background, a linear Ridge regression was trained on the CNN feature representations of all stimuli containing that specific object or background, respectively. The trained decoder was then tested on either the same or every other left-out object or background, resulting in generalization MAE scores between pairs of objects or backgrounds, respectively. This entire procedure was repeated twice using the two instances of each stimulus type that differed only in the views and locations where the objects were placed (see Experimental design). Across these two folds of the cross- validation, one instance of the stimulus was used for training and the second one was used for testing. Finally, we averaged the MAE scores obtained for each pair of objects or backgrounds across the two folds to derive the cross-validated pairwise pattern of generalization performance scores for the model layer. This scheme allowed us to test the prediction performance when testing on the same condition in exactly the same way as when testing on a different condition.

### Mapping low-level statistics to the numerosity predictions

To assess the extent to which the selected low-level statistics could account for the networks’ numerosity predictions across stimuli, we used multiple linear regression with the six image statistics together as predictors and the predicted log-numerosity from the fine-grained generalization across objects or backgrounds, respectively, as the target. We performed this analysis for all the different training conditions of the fine-grained generalization scheme (20 objects & 20 backgrounds). This corresponds to fitting a specific multiple linear regression on the centered 20 x 2 x 1280 = 51 200 log-numerosity values predicted for the test images (19 objects/backgrounds, and the version of the same object/background not seen during training) and the standardized low-level statistics values of these images concatenated together. For each trained object or background, the explained variance (R²) of the regression model was computed, indicating how well the image statistics accounted for the variability in the predicted log- numerosity. We then averaged the explained variance across all the 20 fits, yielding the final average explained variance score for the generalization across objects and backgrounds, respectively.

### Quantifying non-numerical quantity biases on numerosity predictions

To test the influence of non-numerical quantities on numerosity predictions, we performed analyses similar to the ones of (DeWind et al., 2015) whose approach to varying information related to size and spacing we adopted in our stimulus creation. These analyses were performed for the numerosity predictions for test on the same object in the fine-grained generalization scheme (given that quantities as individual object area and perimeter are precisely defined and consistent for the same object but less for objects of different shapes, since the stimulus creation only equated the square area encompassing the object). For each one of the 20 objects, we used the 2560 model predictions for numerosity on the test data as input to a multiple linear regression with Numerosity, Size in Area and Spacing as regressors. The three regressors were standardized and the numerosity predictions were centered. For each network we obtained 20 estimates of (βN, βSzA, βSp) corresponding to the individual objects, of which we subsequently computed their means and standard deviations across objects.

We also used the estimated beta weights for the three regressors (β_N_, β_SzA_, β_Sp_) to compute the angle between this estimated vector and all the axes representing the individual non-numerical dimensions (TA, IA, Spar, FA, TP, Cov and AC) and the three main axes (N, SzA and Sp). In addition, we performed the same analysis with shuffled targets for 10 000 iterations, to simulate the distribution of angles obtained “at chance”. From this distribution, we computed the 95% confidence interval (CI) for which we cannot differentiate the angles lying within this CI from angles obtained through a random procedure.

### Analyses of numerosity selective neurons

To determine whether the results obtained for our photorealistic stimuli were based on the same type of units as observed in previous studies (Kim et al., 2021; Nasr et al., 2019), which were tuned to binary dot sets, we first replicated the experimental approach of these studies. Using a Python translation of the code provided by (Kim et al., 2021), we generated a dataset of visual dot arrays, consisting of images of size 224 × 224 pixels with n = 1, 2, 4, 6, . . . 30 dots pasted at random locations, further split into three different subsets controlling for different non-numerical geometric cues. In the first subset (standard condition), all the dots have nearly identical radii, drawn from the normal distribution N (µ = 7, σ = 0.7). In the second subset (control set 1), the total area covered by the dots remains constant across numerosities (1200 pixels), and the average distance between neighboring dots is constrained within a narrow range (90 to 100 pixels). In the third subset (control set 2), the convex hull formed by the dots is fixed (a pentagon with a circumference of 647 pixels) for numerosities greater than 4, and the individual dots vary randomly in shape between circle, rectangle, ellipse, and triangle.

The preferred numerosity (PN) of an artificial neuron was defined as the numerosity that elicited the largest response on average among the responses for all dot array stimuli. To determine the average tuning curves of all number neurons, the tuning curves of all units were normalized between 0 and 1 and then averaged across neurons with the same PN value.

We identified units as number neurons or non-selective neurons using two-way ANOVA procedures and thresholds as those used in the previous studies: Network units that showed a significant main effect of numerosity (p < 0.01), but no significant effect of stimulus set or interaction between factors (p > 0.01) were labeled as number neurons. Conversely, units that showed a significant main effect of stimulus set (p < 0.01) but no significant effect of numerosity or the interaction between factors (p > 0.01) were labeled as non-selective neurons. Because it has been suggested that the detection of number neurons may be unreliable with small ANOVA sample sizes (Zhang & Wu, 2020), we used 900 images for each numerosity and stimulus set, resulting in a total of 900 × 16 × 3 = 43 200 images, comparable to the number of units in the network layer analyzed (AlexNet conv5).

Finally, we evaluated the decoding performance based on the number neurons compared to the non-selective neurons in the coarse-grained generalization analysis on our dataset of photorealistic stimuli. For comparison, and to assess the extent to which the numerosity information in the CNN representations is only carried by the units defined as number neurons, we also performed this analysis with the entire population of artificial units, but without the numerosity-selective ones. Unlike our previous analyses, the stimuli from both subitizing and estimation ranges were combined in this analysis to decode numerosity N ∈ {1, 2, 4, 6, 10, 24}. In addition, to further evaluate whether the number of units used for the decoding affected numerosity prediction performance, we performed the same decoding analysis by randomly sampling different fractions of the total artificial unit population. For each percentage p of the population retained for the decoding, we bootstrapped the procedure ⌊100/p⌋ + 1 times, allowing for statistical sampling of the entire population of units across the iterations.

Complementary analyses were conducted defining numerosity selective units on our own stimuli, for both the original photorealistic images and their binarized counterparts. Separately for each stimulus type (photorealistic, binarized), numerical range (subitizing, estimation), and stimulus version (independent halves of the dataset differing only by randomly drawn positions and views) we preformed three-way ANOVAs with factors of numerosity (4 levels), size in area (4 levels) and spacing (4 levels). Units were considered as numerosity selective if they had a significant main effect of numerosity (p < .01) and no significant main effect of or interaction with one or both of the other factors (p > .01). For each set of selective units as well as the rest of the population excluding the selective ones, decoding performance in the coarse-grained generalization analysis and the fine-grained generalization analysis across objects was subsequently evaluated on the independent half of the dataset not used for unit definition, to avoid selection bias.

### Statistical analysis

All statistical variables, including the sample sizes, exact p-values, and statistical methods, are reported in the corresponding texts or figure legends. Repeated-measures ANOVA and two-tailed Wilcoxon signed-rank tests were used for analyses comparing the predictive performance between different models. The Wilcoxon signed-rank tests correspond to the paired version of this test, computed by first taking the difference between trained and untrained predictive MAE (corresponding to a 15-sample test for Fig 3 and 30 samples for Fig 5). Pearson’s r was used for all reported correlations.

## Author contributions

Conceptualization: EE, TC,

BT Methodology: TC, EE, BT

Investigation: TC

Visualization: TC

Supervision: EE, BT

Writing—original draft: TC, EE

Writing—review and editing: TC, EE, BT

## Data and materials availability

The image data set and code used to generate the results will be made publicly available in an Open Science Framework repository at https://osf.io/km23y/ upon publication.

## Supporting information

**Fig S1.**
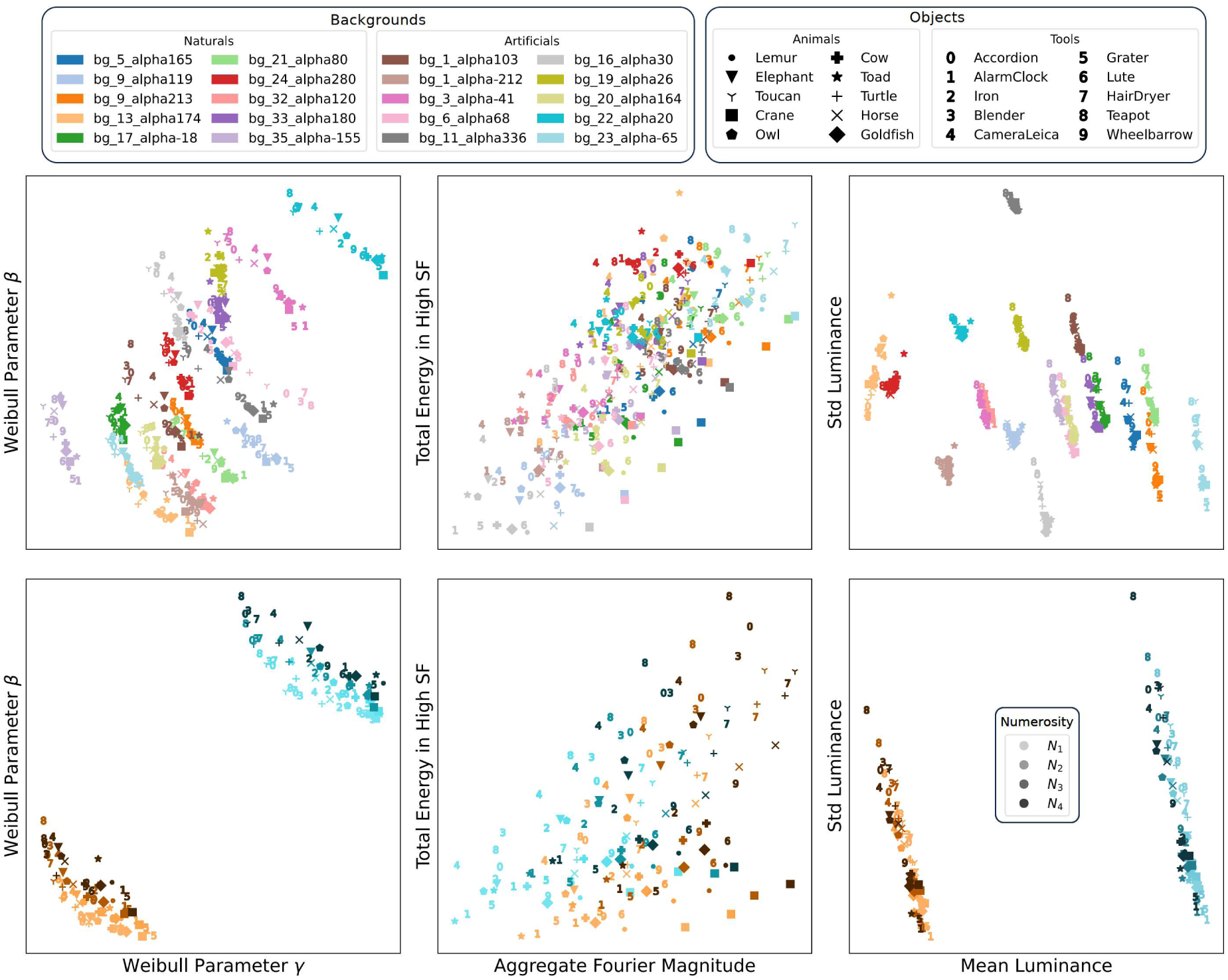
Visualization of the characteristics of the photorealistic stimulus dataset in the space of low-level summary statistics of the images - subitizing range: 1, 2, 3 or 4 objects. (Top row) Overview of the low-level statistics for every one of the 400 (object, background) pairs, depicted by a specific pair of (marker, color). Each plotted point represents the position of a given condition (object, background) averaged across the remaining parametric space dimensions (N, SzA, Sp) and the two versions of each stimulus (which vary only by the random locations of the objects) accounting for a total of 4x4x4x2=128 stimuli averaged. (Bottom row) Overview of the low-level statistics as a function of numerosity for two selected backgrounds. Each plotted point represents the position of a specific condition (numerosity, object) averaged across the remaining non-numerical dimensions (SzA, Sp) and the two versions of each stimulus, accounting for a total of 4x4x2=32 stimuli averaged.

**Fig S2.**
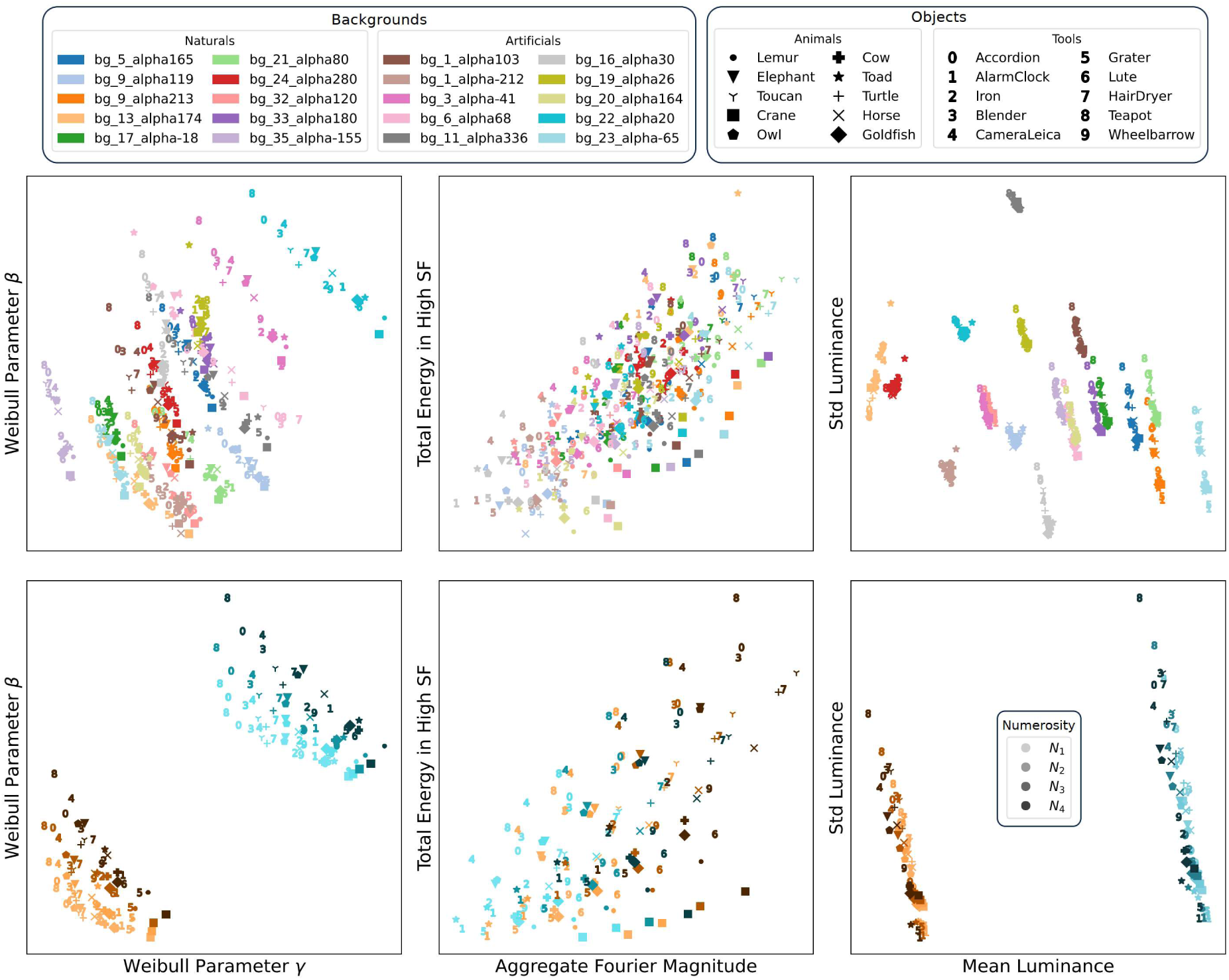
Visualization of the characteristics of the photorealistic stimulus dataset in the space of low-level summary statistics of the images - estimation range: 6, 10, 15 or 24 objects. (Top row) Overview of the low-level statistics for every one of the 400 (object, background) pairs, depicted by a specific pair of (marker, color). Each plotted point represents the position of a given condition (object, background) averaged across the remaining parametric space dimensions (N, SzA, Sp) and the two versions of each stimulus (which vary only by the random locations of the objects) accounting for a total of 4x4x4x2=128 stimuli averaged. (Bottom row) Overview of the low-level statistics as a function of numerosity for two selected backgrounds. Each plotted point represents the position of a specific condition (numerosity, object) averaged across the remaining non-numerical dimension (SzA, Sp) and the two versions of each stimulus, accounting for a total of 4x4x2=32 stimuli averaged.

**Fig S3.**
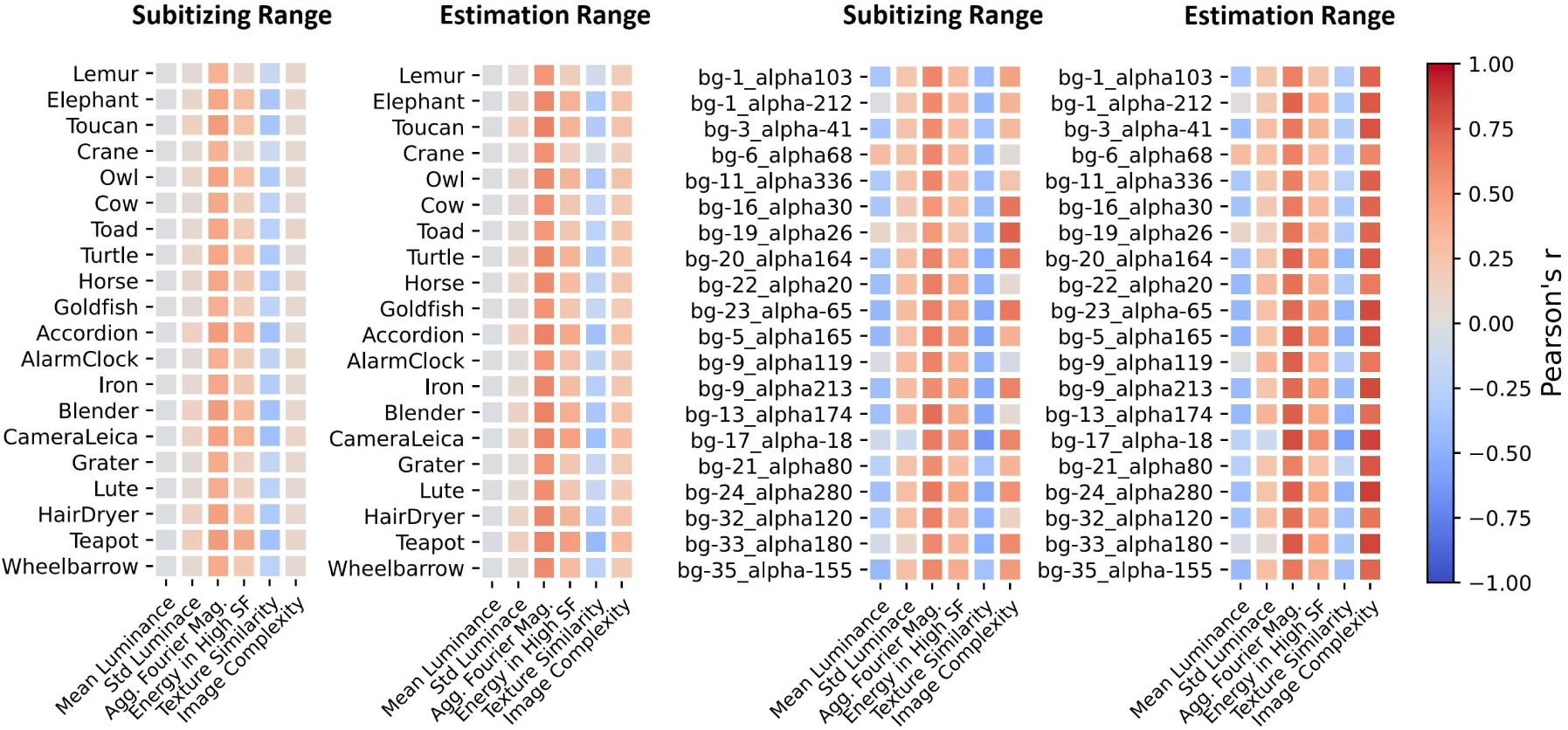
Correlation between log-numerosity and low-level statistics per object/background. Summary of the Pearson correlation between log-numerosity and the six quantified low-level image statistics within our photorealistic stimulus datasets, computed for both numerosity ranges, subitizing (1, 2, 3 or 4 objects) and estimation (6, 10, 15 or 24 objects), across all stimuli sharing the same object (left two subplots) or background (right two subplots), respectively.

**Fig S4.**
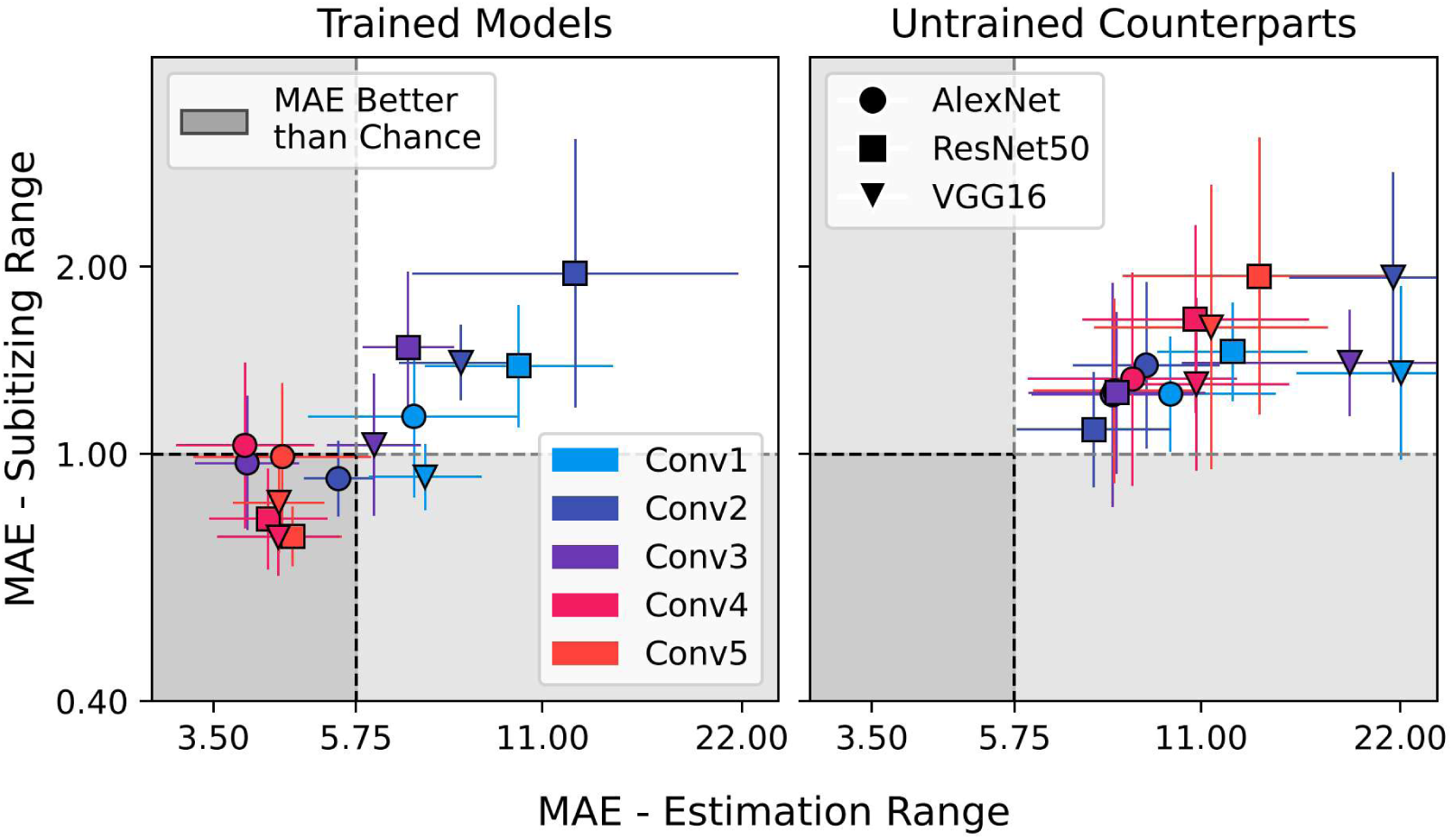
Decoding of numerosity with linear labels. Numerosity prediction performance (mean absolute error, MAE) in the coarse-grained generalization scheme for the different CNN architectures (markers) and layers (colors), when using numerosity itself (rather than the logarithm of numerosity) as the target for Ridge regression. Perfect performance is 0 and simulated chance performance is indicated by a dashed line. Error bars represent STD of the MAE over all the train-test combinations and cross-validation iterations. The subitizing range (y-axis) corresponds to 1 to 4 objects whereas the estimation range (x-axis) corresponds to 6 to 24 objects.

**Fig S5.**
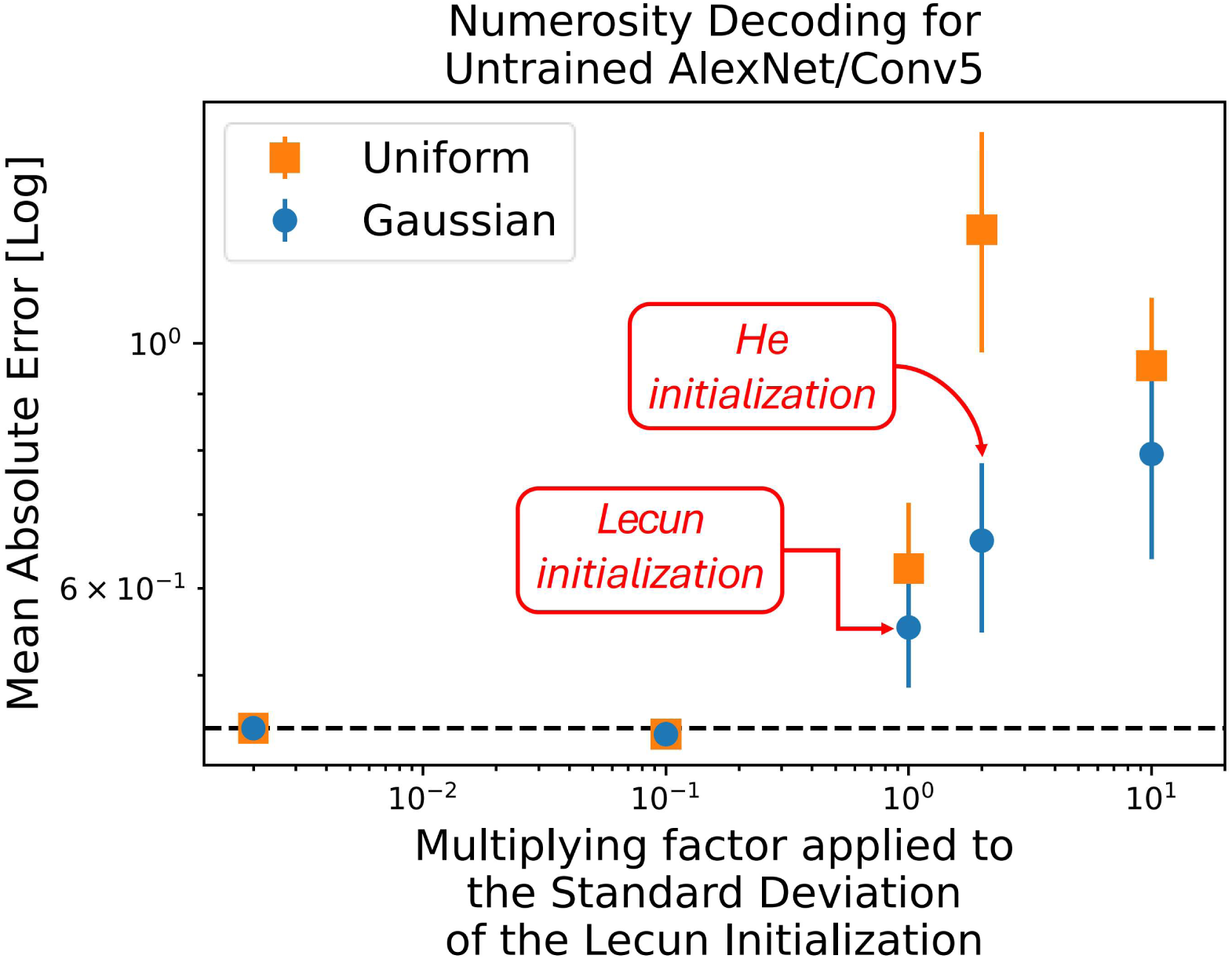
Effect of random initialization of network weights on numerosity decoding in untrained networks. Decoding performance (MAE) is shown for the fifth layer of the untrained version of AlexNet (Random AlexNet - Conv5) for numerosity decoding in the estimation range in the coarse-grained generalization scheme. The different markers represent different hyperparameters of the random initialization (uniform vs Gaussian distribution with different standard deviations). Perfect performance is 0 and simulated chance performance is indicated by a dashed line. Error bars represent STD of the MAE over all the train-test combinations and cross-validation iterations.

**Fig S6.**
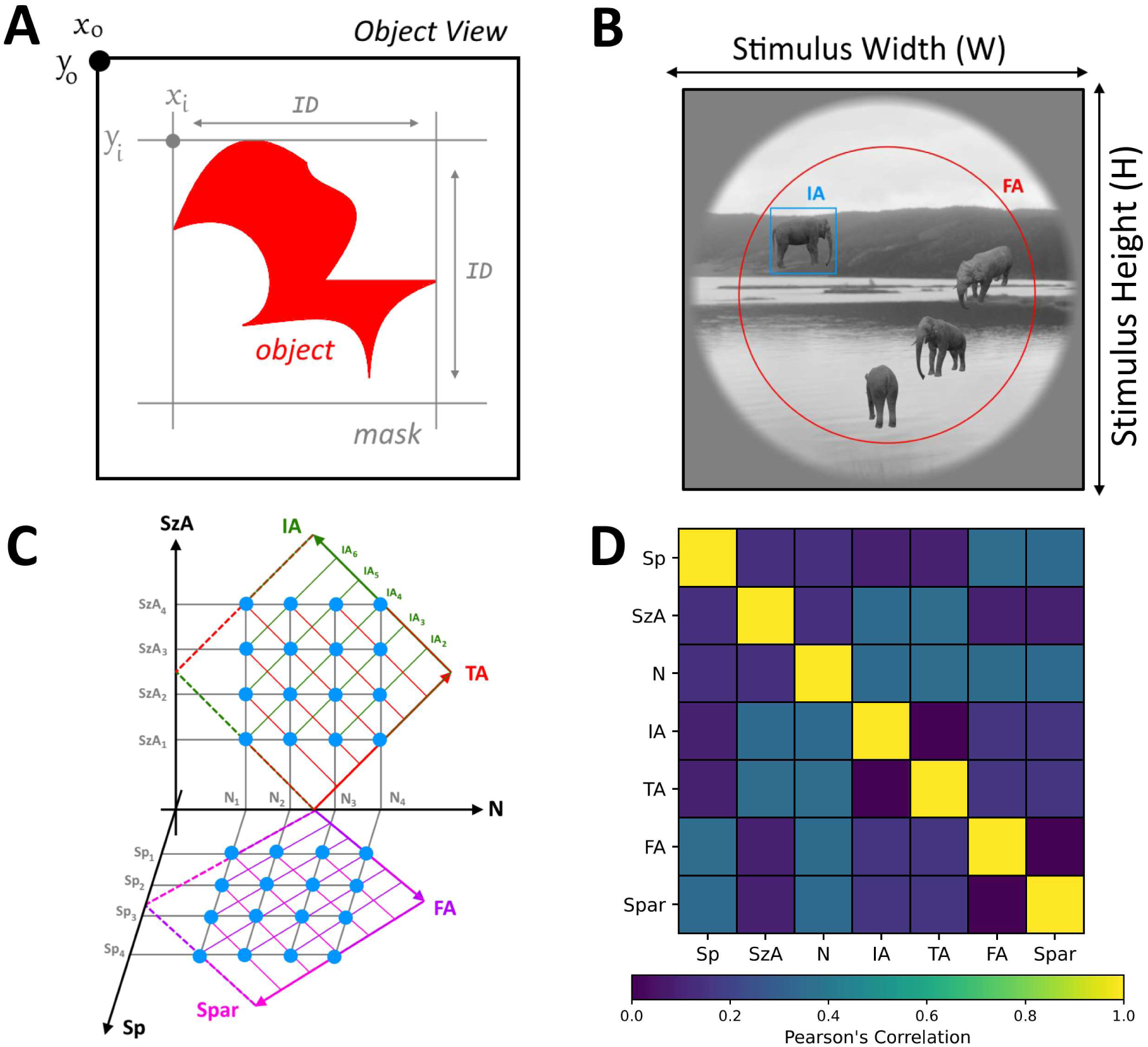
Strategy to control non-numerical geometric stimulus properties in the stimulus dataset. Our stimulus design used a logic following De Wind et al., 2015, Cognition, for creation of multi-item patterns, adapted here to diverse object shapes rather than dots. **(A)** Definition of Item Diameter (ID): The 48 distinct views of a single object do not share an identical “physical” area, corresponding to the number of pixels encompassed by the object’s shape (depicted in red). For the stimulus generation, we considered as the item diameter (ID) of an object the minimal size of the square (illustrated in grey) that encloses the shape of the object, and as item area (IA) the area of this square. **(B)** Stimulus example illustrating the field area (FA) corresponding to the circle within which the objects were allowed to fall, and the IA of one object. **(C)** The two dimensions “Spacing” (Sp, corresponding to an equal relative increase in field area FA and sparsity Spar) and “Size in Area” (SzA, corresponding to an equal relative increase in item area IA and total area TA) were varied orthogonally to numerosity N in four equidistant steps on a log scale. As a result, the constructs Sp and SzA are decorrelated from N whereas the individual non-numerical parameters (IA, TA, FA, Spar) are equi-correlated with N as can be seen in **(D)**.

**Fig S7.**
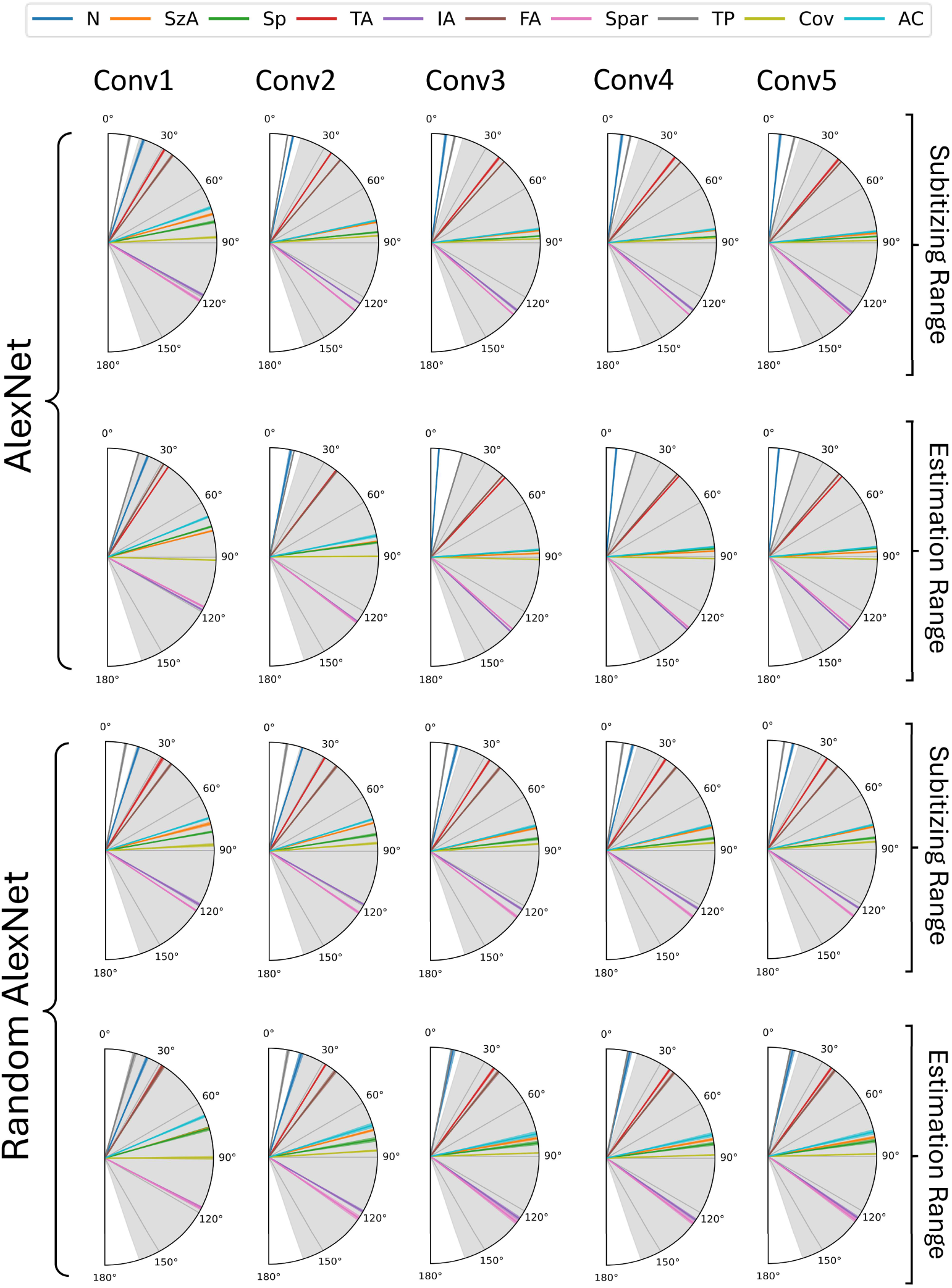
Effects of numerosity and non-numerical quantities on numerosity predictions. Results of quantifying the angle between the vector of weights (β_N_, β_SzA_, β_Sp_) estimated in a multiple regression on numerosity predictions (for test on the same object in the fine-grained decoding scheme) and the different quantities corresponding to directions in the parametric stimulus space. The dark colored lines correspond to the mean and the lighter colored surround to the STD of angles obtained across the 20 different objects. The grey region corresponds to the 95% confidence interval (CI) of angles obtained through a random procedure (in analyses performed with shuffled targets for 10 000 iterations).

**Fig S8.**
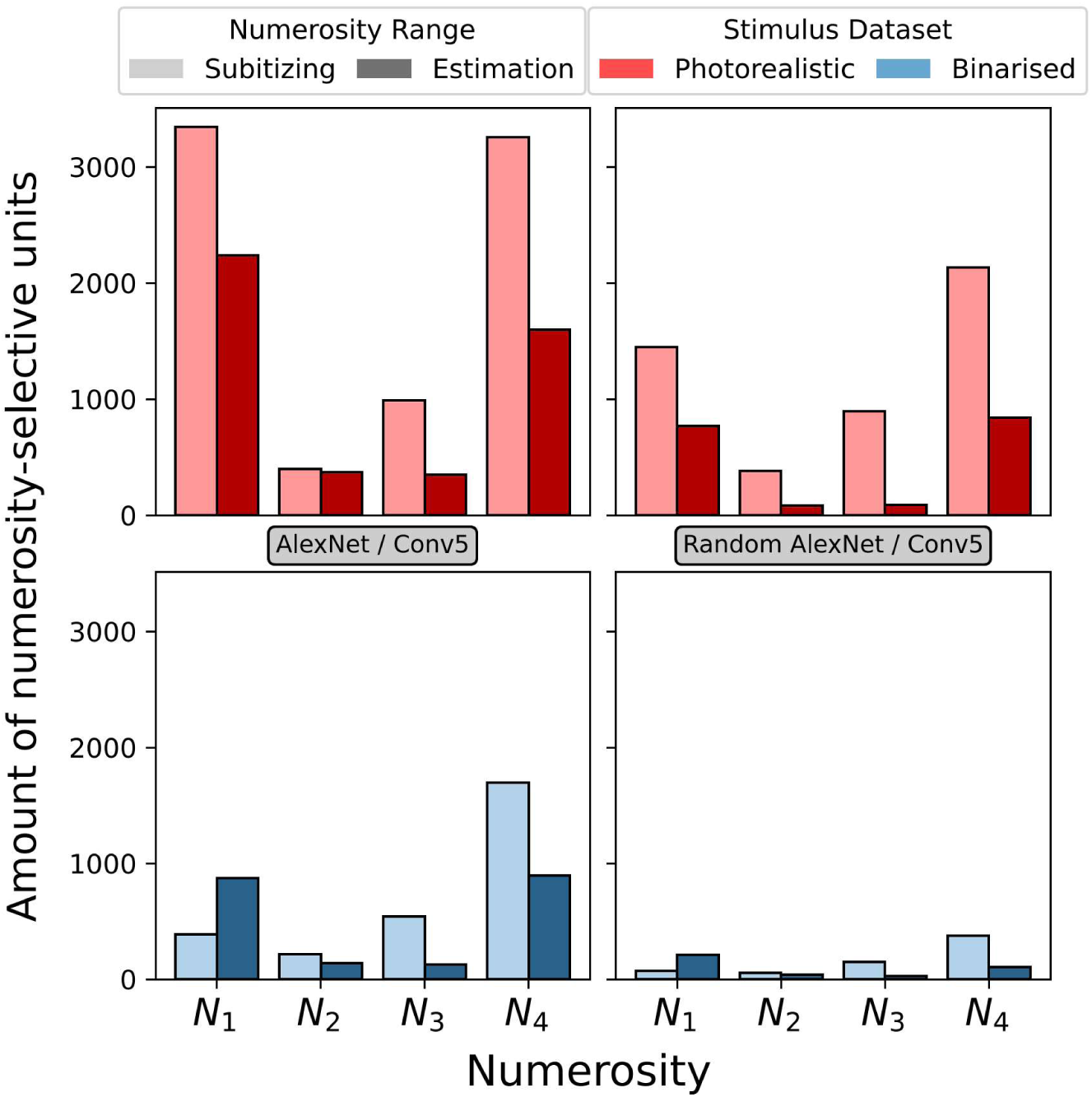
Numerosity selective units for definition on current stimulus datasets. Selective units in layer conv5 of trained and untrained AlexNet were defined as having a main effect of numerosity, and no main effect of or interaction with SzA or Sp, separately for numerical ranges and stimulus versions, for the original photorealistic images of the present dataset as well as their binarized versions. Bar plots indicate of the amounts of selective units (averaged across stimulus versions) for each individual numerosity within each numerical range (N1 to N4 for subitizing: 1 to 4 objects, estimation: 4 to 24 objects) and dataset type (photorealistic, binarized).

**Fig S9.**
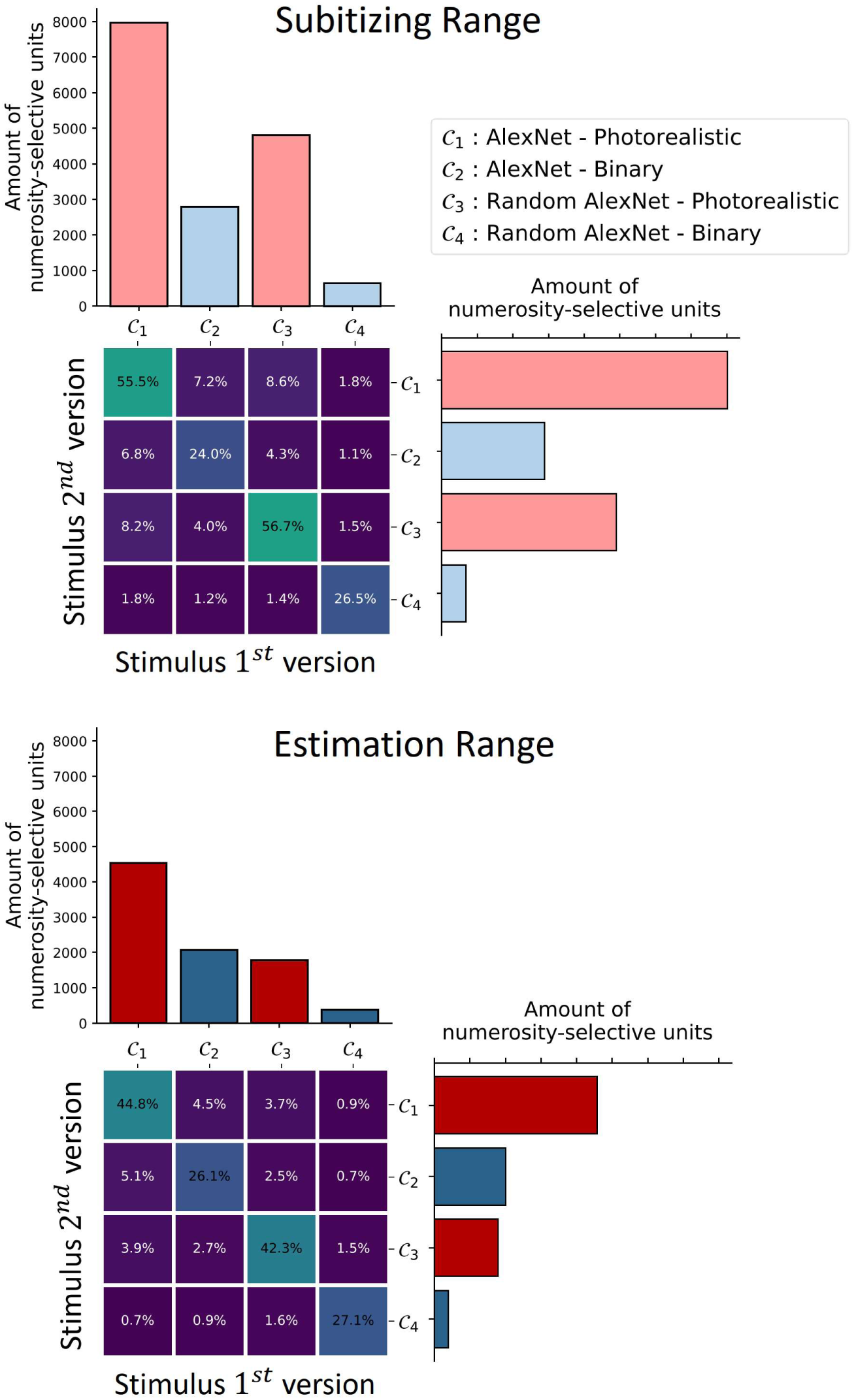
Numerosity selective units and their overlap. Selective units in layer conv5 of trained and untrained AlexNet were defined as having a main effect of numerosity, and no main effect of or interaction with SzA or Sp, separately for numerical ranges and stimulus versions, for the original photorealistic images of the present dataset as well as their binarized versions. Bar plots indicate the total numbers of selective units as defined by these criteria, for the first and second version of the stimuli (independent halves differing by the randomly selected positions and views of the object). Matrices give the percentage of overlap of these units across stimulus versions, dataset types (photorealistic, binarized) and trained vs untrained versions of the network. Percent overlap is computed for a given pair as: intersection of selective units’ indices / union of selective unit indices * 100.

**Fig S10.**
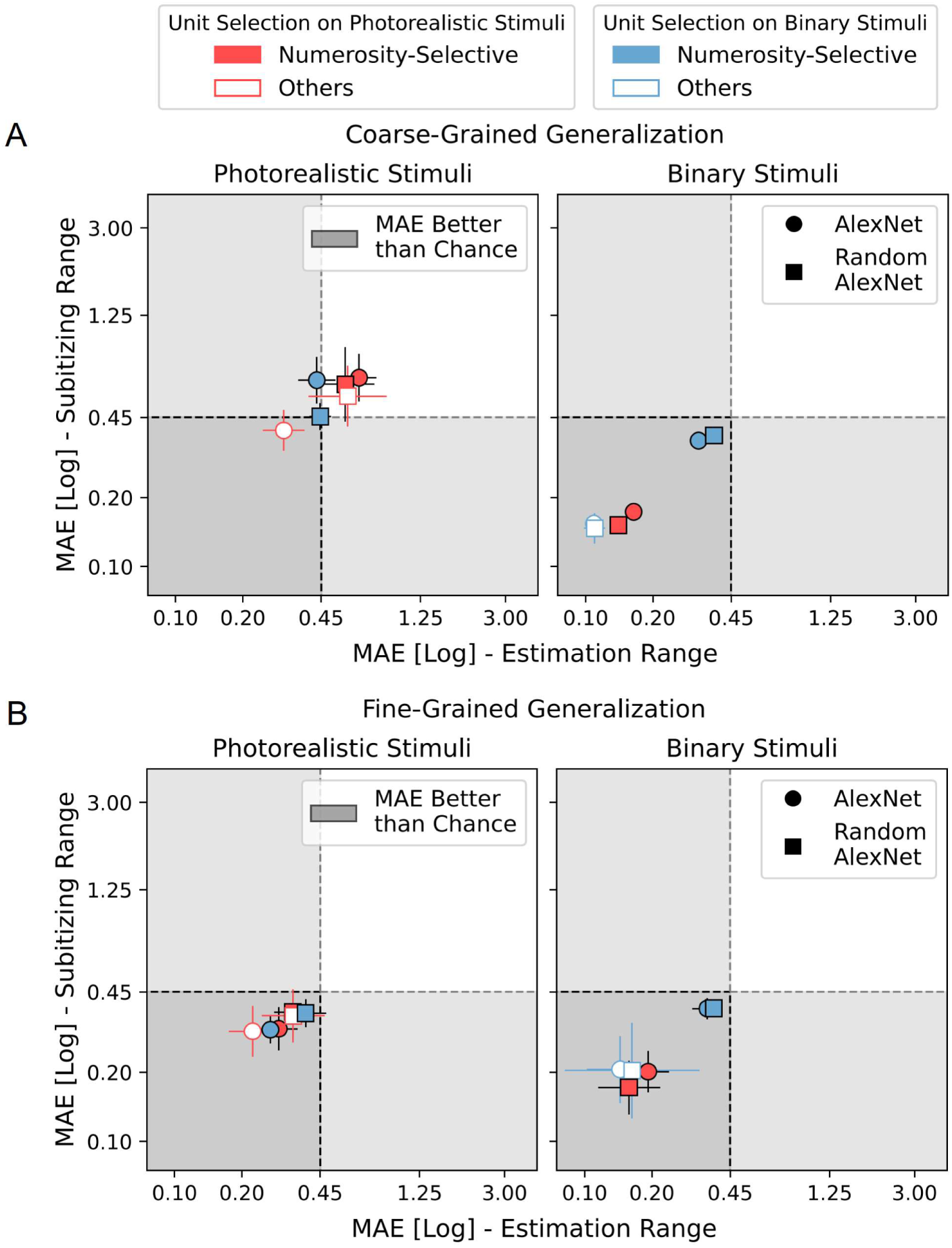
Numerosity decoding for selective units and the rest of the population. Numerosity-Selective units were defined as having a main effect of numerosity, and no main effect of or interaction with SzA and/or Sp, separately for numerical ranges and stimulus versions, either for the original images of the present dataset or their binarized versions. Others units correspond to the larger population excluding the numerosity-selective ones. Decoding was performed on the version of the dataset not used for selectivity definition, to avoid selection bias. Results shown are for layer Conv5 of trained and untrained AlexNet. A) Numerosity decoding performance (MAE) for the coarse-grained generalization scheme performed on either photorealistic or binarized stimuli. B) Numerosity decoding performance (MAE) for the fine- grained generalization scheme across objects performed on either photorealistic or binarized stimuli. Perfect performance is 0 and simulated chance performance is indicated by a dashed line. Error bars represent STD of the MAE over all the train-test combinations and cross- validation iterations.

**S1 Table.**
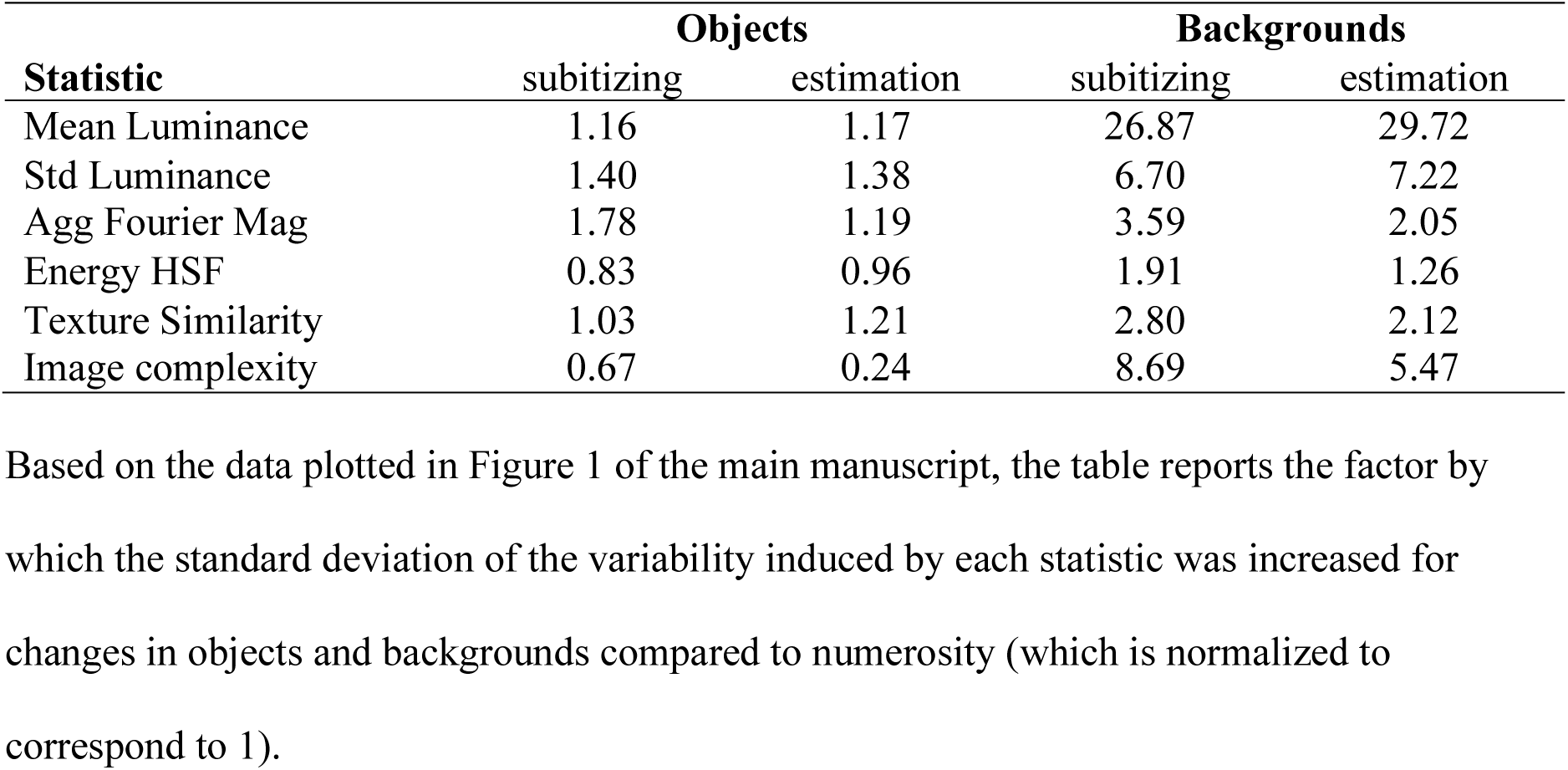
Quantification of degree of variation in low-level summary statistics.

| **Statistic** | **Objects** |  | **Backgrounds** |  |
| --- | --- | --- | --- | --- |
|  | subitizing | estimation | subitizing | estimation |
| Mean Luminance | 1.16 | 1.17 | 26.87 | 29.72 |
| Std Luminance | 1.40 | 1.38 | 6.70 | 7.22 |
| Agg Fourier Mag | 1.78 | 1.19 | 3.59 | 2.05 |
| Energy HSF | 0.83 | 0.96 | 1.91 | 1.26 |
| Texture Similarity | 1.03 | 1.21 | 2.80 | 2.12 |
| Image complexity | 0.67 | 0.24 | 8.69 | 5.47 |

**S2 Table.**
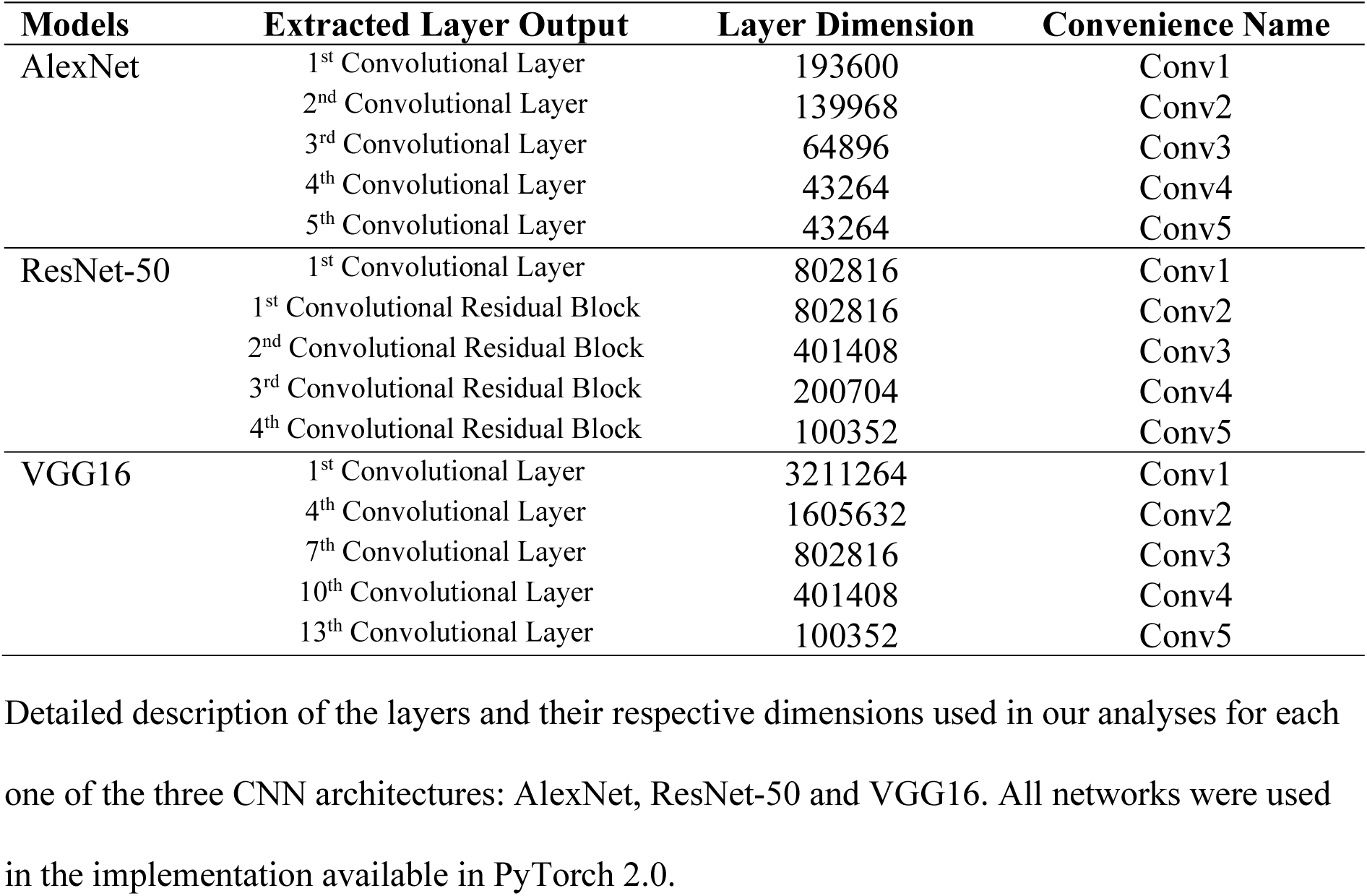
Hierarchical convolutional networks and layers investigated.

**S3 Table.** Most predictive layers in fine-grained generalization.

| Network | Objects |  | Backgrounds |  |
| --- | --- | --- | --- | --- |
|  | Test same | Test diff | Test same | Test diff |
| AlexNet trained | 4 / 4 | 5 / 4 | 4 / 5 | 4 / 5 |
| ResNet-50 trained | 4 / 4 | 4 / 4 | 4 / 4 | 5 / 5 |
| VGG16 trained | 5 / 5 | 5 / 3 | 5 / 5 | 5 / 5 |
| AlexNet untrained | 3 / 5 | 5 / 5 | 5 / 5 | 4 / 2 |
| ResNet-50 untrained | 4 / 4 | 4 / 4 | 4 / 4 | 3 / 5 |
| VGG16 untrained | 5 / 5 | 2 / 2 | 5 / 5 | 5 / 5 |

